# Functional diversification gave rise to allelic specialization in a rice NLR immune receptor pair

**DOI:** 10.1101/2021.06.25.449940

**Authors:** Juan Carlos De la Concepcion, Javier Vega Benjumea, Aleksandra Białas, Ryohei Terauchi, Sophien Kamoun, Mark J. Banfield

## Abstract

Cooperation between receptors from the NLR superfamily is important for intracellular activation of immune responses. NLRs can function in pairs that, upon pathogen recognition, trigger hypersensitive cell death and stop pathogen invasion. Natural selection drives specialization of host immune receptors towards an optimal response, whilst keeping a tight regulation of immunity in the absence of pathogens. However, the molecular basis of co-adaptation and specialization between paired NLRs remains largely unknown. Here, we describe functional specialization in alleles of the rice NLR pair Pik that confers resistance to strains of the blast fungus *Magnaporthe oryzae* harbouring AVR-Pik effectors. We revealed that matching pairs of allelic Pik NLRs mount effective immune responses whereas mismatched pairs lead to autoimmune phenotypes, a hallmark of hybrid necrosis in both natural and domesticated plant populations. We further showed that allelic specialization is largely underpinned by a single amino acid polymorphism that determines preferential association between matching pairs of Pik NLRs. These results provide a framework for how functionally linked immune receptors undergo co-adaptation to provide an effective and regulated immune response against pathogens. Understanding the molecular constraints that shape paired NLR evolution has implications beyond plant immunity given that hybrid necrosis can drive reproductive isolation.

## Introduction

Pathogens use an array of molecules, termed effectors, to successfully colonize hosts (Win et al., 2012). Intracellular detection of effectors relies on immune receptors from the nucleotide-binding, leucine-rich repeats (NLR) superfamily (Bentham et al., 2020; Jones et al., 2016; Saur et al., 2020). Upon recognition, NLRs act as nucleotide-operated switches, exchanging ADP for ATP (Bernoux et al., 2016; Tameling et al., 2002; Wang et al., 2019b; Williams et al., 2011), and oligomerise into supramolecular signalling platforms (Hu et al., 2015; Ma et al., 2020; Martin et al., 2020; Sharif et al., 2019; Tenthorey et al., 2017; Wang et al., 2019a; Zhang et al., 2015). This leads to immune responses, including programmed cell death, that restrict pathogen growth. The assembly of such sophisticated molecular machinery needs to be well coordinated and tightly regulated to ensure an efficient immune response, while avoiding the deleterious effect of constitutive immune activation (Chae et al., 2016; Karasov et al., 2017; Li et al., 2020; Richard and Takken, 2017).

NLRs form the most expanded and diversified protein family in plants (Meyers et al., 2003; Van de Weyer et al., 2019; Yue et al., 2012). Since their discovery, plant NLRs have been heavily studied and around 450 NLR proteins from 31 genera of flowering plants have been functionally validated (Kourelis et al., 2021). Plant NLRs present multiple layers of complexity (Barragan and Weigel, 2020), often functioning in genetically linked pairs (Eitas and Dangl, 2010; Griebel et al., 2014) or as part of complex immune networks (Wu et al., 2018). In such cases, NLRs specialize their role in immune activation, acting as “sensors” that detect pathogen effectors or as “helpers” that amplify and propagate immune signalling (Adachi et al., 2019b; Jubic et al., 2019). Paired NLRs are prevalent in plant genomes (Stein et al., 2018; Wang et al., 2019c) with a subset of sensor NLRs harbouring atypical domains integrated into their architecture (Bailey et al., 2018; Kourelis et al., 2021; Kroj et al., 2016; Sarris et al., 2016). These domains can be derived from pathogen host targets that act as sensor domains within NLRs by binding pathogen effectors (Bialas et al., 2018; Cesari et al., 2014a; Maidment et al., 2021; Oikawa et al., 2020).

Cooperating NLRs must balance a trade-off between adaptive evolution to fast evolving pathogens and maintaining a fine-tuned regulation of complex receptor assemblies. NLRs with different evolutionary trajectories may drift apart and eventually mismatch. When these mismatched NLRs are combined in the same individual through genetic crossing, constitutive immune activation can occur leading to deleterious phenotypes including dwarfism, necrosis, and lethality (Bomblies et al., 2007; Chae et al., 2014). These “Dangerous Mix” phenotypes are known in plant breeding as hybrid necrosis and have important implications in agriculture (Caldwell and Compton, 1943; Calvo-Baltanás et al., 2021; Hermsen, 1963a, b; Li and Weigel, 2021; Wan et al., 2021; Yamamoto et al., 2010). In Arabidopsis, two genetically unlinked NLR proteins encoded on different chromosomes were shown to physically associate in the mixed immune background of hybrid plants, underpinning hybrid necrosis (Tran et al., 2017). Similarly, association between NLRs and alleles of non-NLR proteins derived from a different genetic background was also shown to induce NLR activation and autoimmune phenotypes (Barragan et al., 2019; Li et al., 2020). However, the biochemical basis of adaptive specialization in genetically linked NLR receptor pairs remains largely unknown. In particular, we know little about how coevolution between paired NLRs has impacted their activities. We lack a validated framework to explain how plant immune receptors adapt and specialize, even though this process has important consequences for plant diversification and evolution (Bomblies and Weigel, 2007; Calvo-Baltanás et al., 2021; Dobzhansky, 1937; Li and Weigel, 2021).

The rice NLRs *Pik-1* and *Pik-2* form a linked gene pair arranged in an inverted configuration on chromosome 11 with only ∼2.5 Kb separating their start codons (Ashikawa et al., 2008). The Pik pair is present in the genetic pool of rice cultivars as two major haplotypes (Bialas et al., 2021; Kanzaki et al., 2012). Pik pairs belonging to the K haplotype confer resistance to strains of the rice blast fungus, *Magnaporthe oryzae*, that harbour the effector AVR-Pik (Ashikawa et al., 2008). The sensor NLR Pik-1 binds AVR-Pik effectors through a heavy metal–associated (HMA) domain integrated into its architecture (De la Concepcion et al., 2018; Kanzaki et al., 2012; Maqbool et al., 2015). Upon effector recognition, Pik-1 cooperates with the helper NLR Pik-2 to activate immune signalling (Zdrzalek et al., 2020) that leads to pathogen resistance. The Pik NLR pair occurs as allelic series in both Japonica and Indica rice cultivars (Chaipanya et al., 2017; Costanzo and Jia, 2010; Hua et al., 2012; Xu et al., 2008). The AVR-Pik effectors are also polymorphic and present signatures of selection (Bentham et al., 2021; Bialas et al., 2018; Yoshida et al., 2009). Allelic Pik NLRs have differential recognition specificities for the AVR-Pik variants (De la Concepcion et al., 2021; Kanzaki et al., 2012), which is underpinned by differential effector binding to the Pik-1 HMA domain (De la Concepcion et al., 2018; De la Concepcion et al., 2021; Maqbool et al., 2015). Two allelic variants of Pik-1, Pikp-1 and Pikm-1, acquired high-affinity binding to the *M. oryzae* AVR-Pik effector through convergent evolution of their HMA domains (Bialas et al., 2021). Additionally, Pikm-1 and Pikh-1 alleles have been shown to convergently evolve towards extended recognition specificity of AVR-Pik variants (De la Concepcion et al., 2021). This adaptive evolution towards recognition of rapidly evolving effectors has led to marked diversification of the integrated HMA domain (Bialas et al., 2021). As a consequence, Pik-1 HMA domain is the most sequence-diverged domain in the Pik NLR pair (Bialas et al., 2021; Bialas et al., 2018; Costanzo and Jia, 2010).

While Pik-1 acts as a sensor, Pik-2 acts as a helper NLR that is required for the activation of immune responses (Maqbool et al., 2015; Zdrzalek et al., 2020). Evolutionary analyses have shown that the genetic linkage of this NLR pair is ancient and revealed marked signatures of adaptive evolution in the integrated HMA domain of Pik-1 (Bialas et al., 2021). However, little is known about sensor/helper coevolution in Pik and how these multidomain proteins have adapted to changes in the rapidly evolving integrated HMA domain of Pik-1.

Here, we used two allelic variants of Pik, Pikp and Pikm, to explore NLR sensor/helper specificity **(Figure S1)**. We challenged the hypothesis that throughout evolutionary time, these two allelic Pik pairs have become diverged and mismatched. Indeed, mismatched pairs of Pik-1 and Pik-2 display constitutive cell death reminiscent of autoimmune phenotypes. We identified a single amino acid polymorphism in the helper NLR Pik-2 that underpins both allelic specialization and immune homeostasis. This finding allowed to reconstruct the evolutionary history of this coevolution. Altogether, these results demonstrate that NLR pairs can undergo co-adaptation and functional specialization, offering a molecular framework to understand how they evolve to respond to pathogen effectors while maintaining a tight regulation of immune responses.

## Results

### A coevolved Pik NLR pair is required for efficient cell death response to AVR-Pik effectors in *N. benthamiana*

Two of the most studied Pik alleles, Pikp (cv. K60) and Pikm (cv. Tsuyuake), fall into phylogenetically distinct groups (Bialas et al., 2021; De la Concepcion et al., 2021; Kanzaki et al., 2012). Pikm originated in the Chinese Japonica cultivar Hokushi Tami (Kiyosawa, 1978) while Pikp originated in the Indica cultivar Pusur in Pakistan (Kiyosawa, 1969). Thus, we hypothesised that these alleles have been exposed to differential selection pressures during domestication of elite cultivars and have undergone distinct evolutionary trajectories.

To test for sensor/helper specificity in allelic Pik pairs, we co-expressed the sensor NLR Pikm-1 with either the helper NLR Pikp-2 or Pikm-2 in *N. benthamiana* and assessed the capacity to trigger a cell death in response to rice blast effector variants AVR-Pik D, E or A (**Figure 1**). As previously reported, Pikm pair mediated a hierarchical cell death response in the order of AVR-PikD > AVR-PikE > AVR-PikA (De la Concepcion et al., 2018). However, the intensity of cell death was lower when Pikm-1 was co-expressed with Pikp-2 instead of Pikm-2 **(Figure 1, Figure S2)**. Protein accumulation of both Pikp-2 and Pikm-2 proteins in planta was similar **(Figure S3)**.

**Figure 1.**
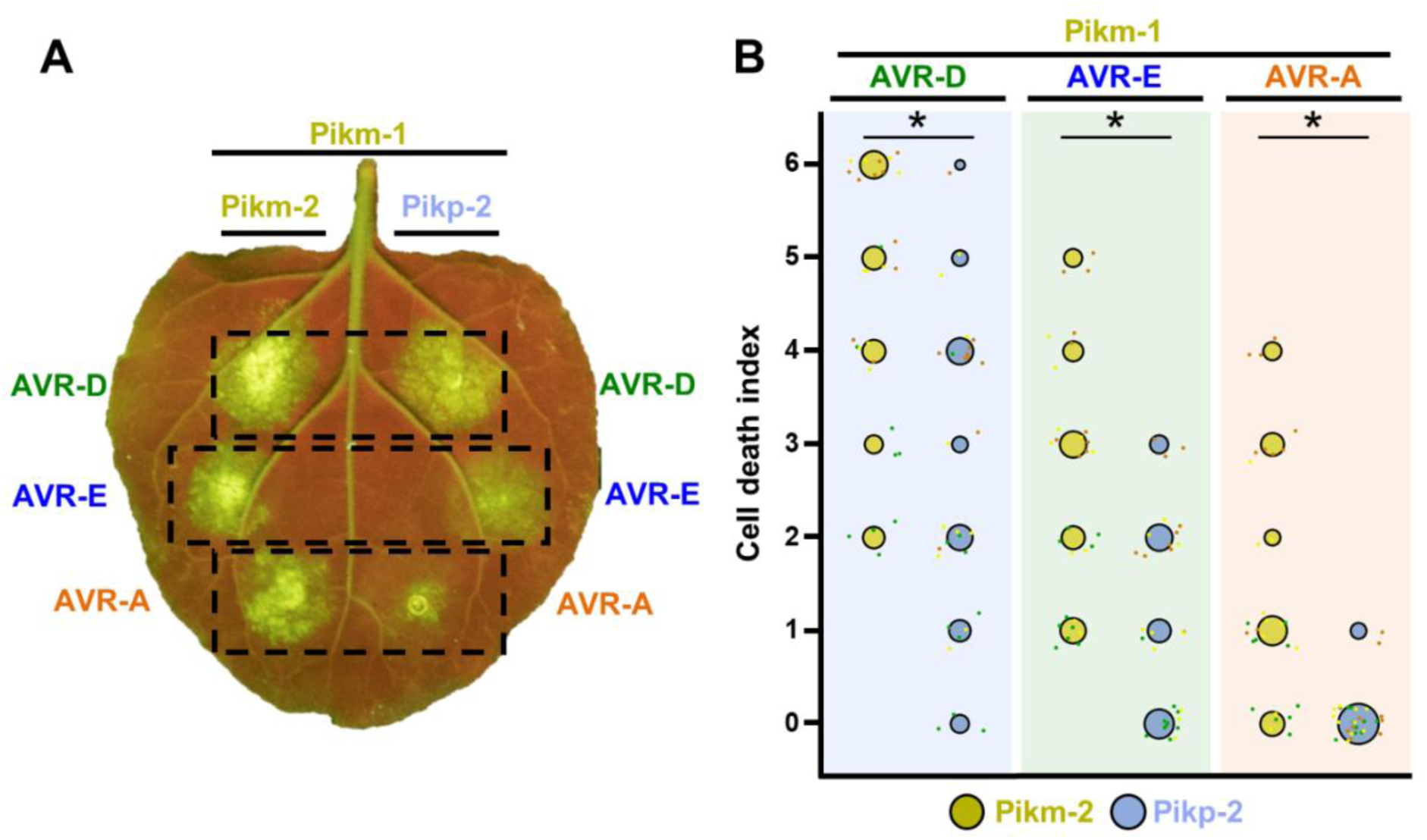
Pikm-1 elicits a stronger response to the AVR-Pik effectors when it is paired with Pikm-2 than with Pikp-2. **(A)** Representative *N. benthamiana* leaf depicting Pik-mediated cell death as autofluorescence under UV light. Pikm-1 was co-expressed with either Pikm-2 or Pikp-2 and the AVR-Pik effector alleles recognized by Pikm. Side-by-side infiltrations are highlighted with dashed boxes. **(B)** Scoring of cell death triggered by Pikp-2 or Pikm-2 with each AVR-PikD (AVR-D), AVR-PikE (AVR-E) and AVR-PikA (AVR-A) is represented as dot plots. The total number of repeats was 30. For each sample, all the data points are represented as dots with a distinct colour for each of the three biological replicates; these dots are jittered around the cell death score for visualisation purposes. The size of the central dot at each cell death value is proportional to the number of replicates of the sample with that score. Significant differences between relevant conditions are marked with an asterisk and the details of the statistical analysis are summarised in **Figure S2**.

These results indicate that Pikm-2 is required for the full Pikm mediated cell death response to the AVR-Pik effectors in *N. benthamiana*. This suggests a possible functional specialization of the helper NLR Pik-2 towards an effective cell death response to these rice blast effectors.

### A single amino acid polymorphism in Pik-2 has an important role in cell death responses to the AVR-Pik effectors

To dissect the basis of the differential cell death phenotypes displayed by Pikp-2 and Pikm-2 in response to the AVR-Pik effectors, we used site-directed mutagenesis to exchange the residues at each of the three Pik-2 polymorphic positions **(Figure S3)**. We then co-expressed Pikm-1 and each of the Pik-2 mutants with either AVR-PikD, AVR-PikE or AVR-PikA. For each assay, we scored the cell death responses and compared the differences with the Pikm control to qualitatively measure the contribution of each polymorphism to cell death **(Figure 2, Figure S4, S5, S6, S7)**. In brief, this assay aimed to identify reciprocal mutations in Pikm-2 and Pikp-2 that may reduce or increase immune responses, when compared with wild-type Pikm-2.

**Figure 2.**
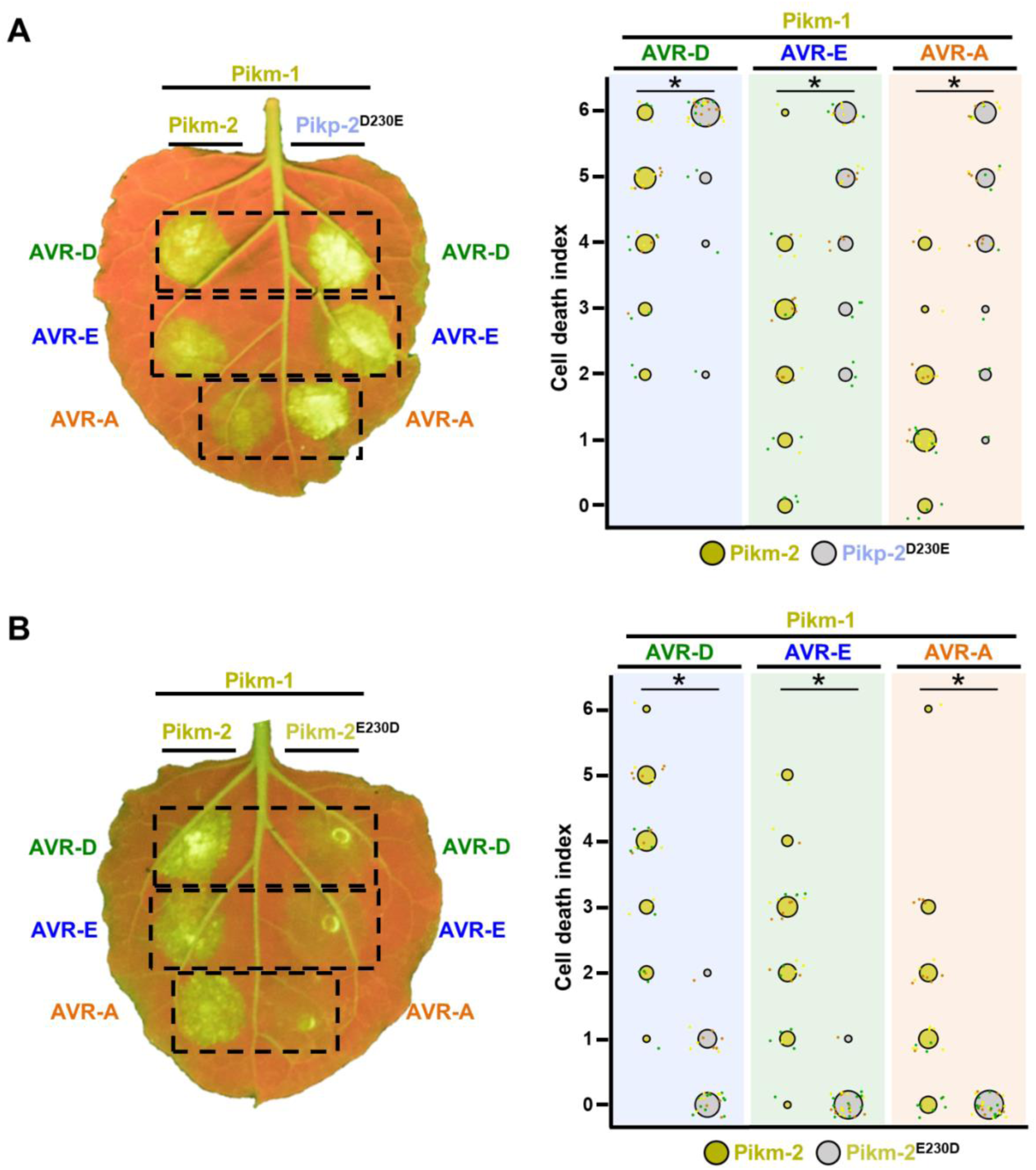
A single Pik-2 polymorphism modulates the cell death response to the AVR-Pik effectors. Representative leaves depicting cell death mediated by Pik-2 mutants as autofluorescence under UV light. Pikm-1 was co-expressed with either **(A)** Pikp-2 Asp230Glu or **(B)** Pikm-2 Glu230Asp and AVR-PikD (AVR-D), AVR-PikE (AVR-E) or AVR-PikA (AVR-A). Side-by-side infiltrations with Pikm NLR pair are highlighted with dashed boxes for comparison. Cell death scoring is represented as dot plots. The number of repeats was 30. For each sample, all the data points are represented as dots with a distinct colour for each of the three biological replicates; these dots are jittered about the cell death score for visualisation purposes. The size of the central dot at each cell death value is proportional to the number of replicates of the sample with that score. Significant differences between relevant conditions are marked with an asterisk and the details of the statistical analysis are summarised in **Figure S4**.

A single amino acid change at position 230 was responsible for the major differences in cell death responses (**Figure 2**). Despite the similar properties of their side chains, the Asp230Glu mutation in Pikp-2 showed an increase in the level of cell death response to AVR-Pik effectors **(Figure 2A, S4A)**. By contrast, the Glu230Asp mutation in Pikm-2 reduced the cell death response to each AVR-Pik effector compared with wild-type Pikm-2, displaying only a slight response to AVR-PikD **(Figure 2B, S4B)**. This points to a major involvement of the Pikm-2 Glu230 residue in the extended response to AVR-Pik effectors observed in Pikm (De la Concepcion et al., 2018).

Mutations at polymorphic positions 434 and 627 did not have the strong effect observed in the mutants at position 230. The Thr434Ser and Met627Val mutations in Pikp-2 did not yield higher levels of cell death response compared with Pikm-2 **(Figure S5A, C, S6A, C, S7A, C)**. Likewise, neither Pikm-2 Ser434Thr nor Pikm-2 Val627Met showed a lower level of cell death response compared with wild-type Pikm-2 **(Figure S5B, D, S6B, D, S7B, D)**. Interestingly, Val627Met in Pikm-2 consistently increased cell death responses, particularly to AVR-PikE and AVR-PikA **(Figure S5D, S6D, S7)** implying a negative contribution of Pikm-2 polymorphism Val627 towards cell death responses. All mutants had a similar level of protein accumulation in *N. benthamiana* compared to wild-type Pikp-2 and Pikm-2 **(Figure S3).**

Altogether, these results demonstrate that polymorphisms in Pik-2 play an important role in facilitating response to different AVR-Pik alleles. Particularly, a single polymorphic residue, Glu230, was revealed as a major determinant of the increased cell death responses to the AVR-Pik effectors displayed by the Pikm NLR pair.

### Mismatched Pik pair Pikp-1/Pikm-2 triggers constitutive cell death responses in *N. benthamiana*

When independently evolved NLR receptors meet in the mixed immune background of a hybrid plant, it can lead to misregulation in the form of suppression (Hurni et al., 2014; Stirnweis et al., 2014) or constitutive activation of immune responses (Chae et al., 2014; Li et al., 2020; Tran et al., 2017).

The Pikp and Pikm allelic pairs trigger a strong cell death response in *N. benthamiana* when co-expressed with rice blast effector AVR-PikD, but not in the absence of effector (De la Concepcion et al., 2018; Maqbool et al., 2015). However, we noticed that when Pikp-1 was co-expressed together with Pikm-2, it led to cell death response in the absence of AVR-PikD (**Figure 3**). We did not observe NLR autoactivation in the reciprocal mismatched pair Pikm-1/Pikp-2 (**Figure 3**).

**Figure 3.**
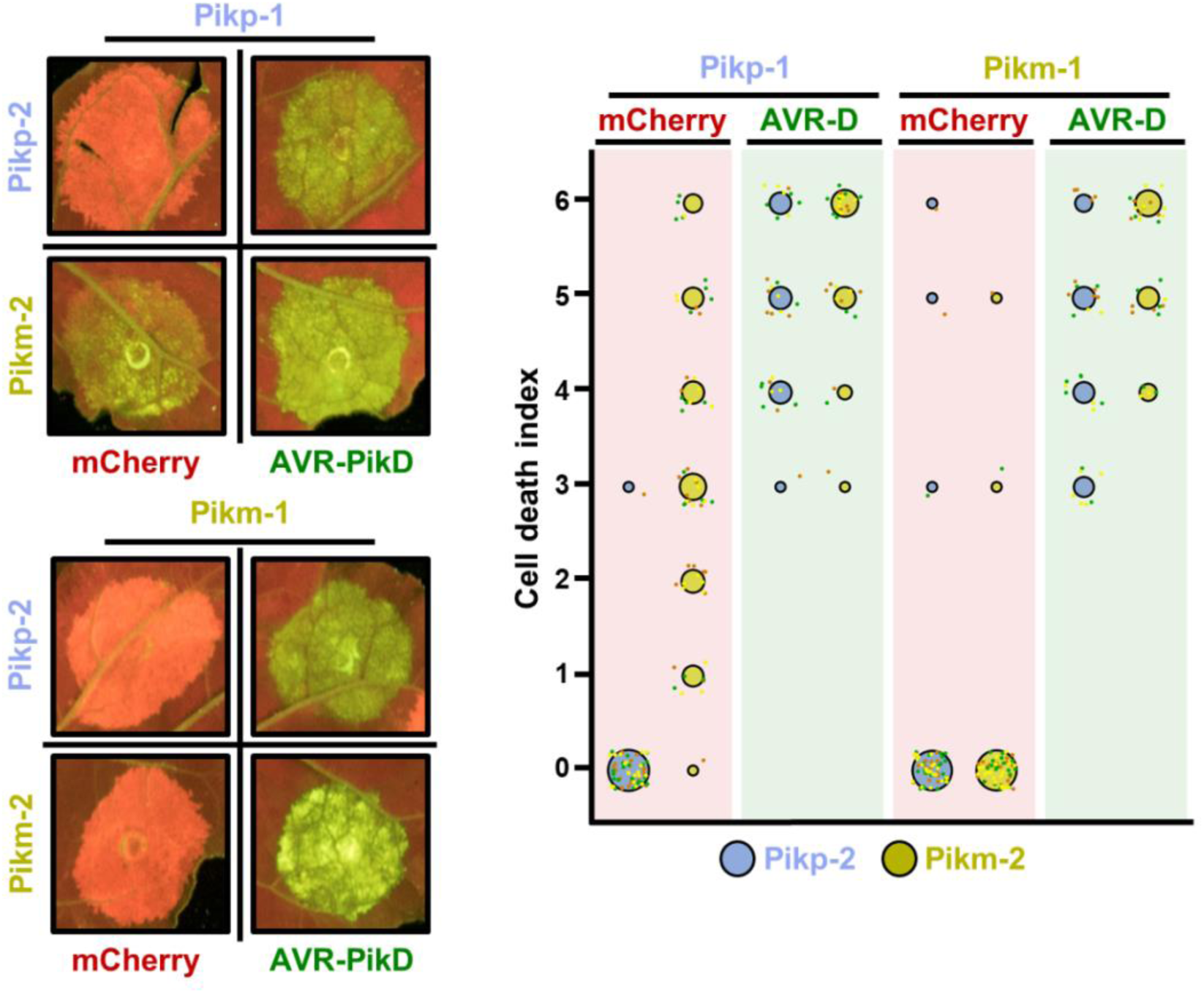
Pikm-2 triggers constitutive cell death in the presence of Pikp-1. Representative leaf spot images and scoring of Pik mediated cell death as autofluorescence under UV-light in the presence or absence of AVR-PikD. Cell death assay scoring represented as dot plots comparing cell death triggered by Pikp-2 and Pikm-2 when co-expressed with Pikp-1 or Pikm-1. The number of repeats was 60 and 30 for the spots co-infiltrated with mCherry and AVR-PikD, respectively. For each sample, all the data points are represented as dots with a distinct colour for each of the three biological replicates; these dots are jittered about the cell death score for visualisation purposes. The size of the central dot at each cell death value is proportional to the number of replicates of the sample with that score.

These results reveal signatures of coevolution in the Pikp and Pikm allelic pairs. We hypothesise that these allelic pairs have coevolved with their respective partners and have drifted enough to trigger a misregulated form of immune response when they are mismatched, leading to constitutive cell death in *N. benthamiana*.

### Pik autoactivity is linked to immune signalling

We sought to gain knowledge on the constitutive cell death mediated by Pikm-2 and understand the link with NLR activation. To this end, we mutated Pikm-2 in the conserved P-loop and MHD motifs and tested their ability to trigger constitutive cell death responses in the absence of the AVR-PikD effector.

The P-loop motif is conserved in NLR proteins and mediates nucleotide binding linked with oligomerization and NLR activation (Ma et al., 2020; Wang et al., 2019b). Loss-of-function mutations at this position render NLRs inactive and have been extensively documented (Tameling et al., 2002; Tameling et al., 2006; Williams et al., 2011). A Lys217Arg mutation in the P-loop motif of Pikp-2 abrogates Pik-mediated cell death responses to AVR-PikD in *N. benthamiana* (Zdrzalek et al., 2020). Introducing this mutation in Pikm-2 abolished Pikm-mediated cell death response to the rice blast effector AVR-PikD (**Figure 4**) and also abrogated the constitutive cell death response triggered by the Pikp-1/Pikm-2 NLR mismatch (**Figure 4**).

**Figure 4.**
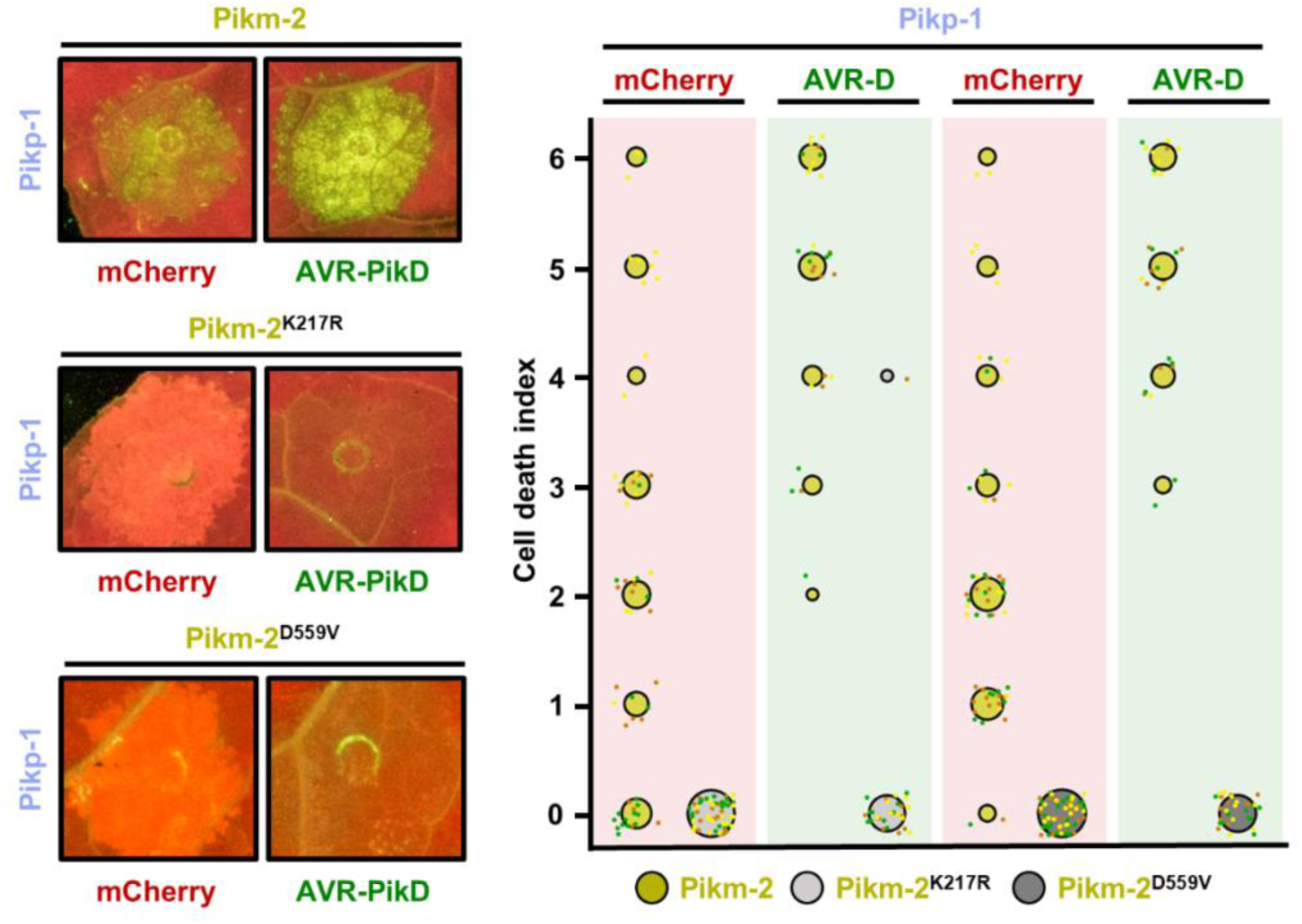
Constitutive cell death in mismatched Pik pairs is dependent on P-loop and MHD motifs. Representative leaf spot images and scoring of Pikm-2 mediated cell death as autofluorescence under UV-light. Cell death scoring is represented as dot plots comparing cell death triggered by Pikm-2 mutant in P-loop (Lys217Arg) and MHD (Asp559Val) motifs and wild-type Pikm-2. Mutants and wild-type proteins were co-expressed with Pikp-1 and mCherry (red panel) or AVR-PikD (green panel). The number of repeats was 60 and 30 for the spots co-infiltrated with mCherry and AVR-PikD, respectively. For each sample, all the data points are represented as dots with a distinct colour for each of the three biological replicates; these dots are jittered about the cell death score for visualisation purposes. The size of the central dot at each cell death value is proportional to the number of replicates of the sample with that score.

NLR activities are also altered by mutations in the MHD motif. An Asp to Val mutation in this motif is predicted to change ATP/ADP binding preference and, in many cases, renders NLRs constitutively active (Bernoux et al., 2016; Tameling et al., 2006; Williams et al., 2011). Contrary to other NLRs, introducing Asp559Val in the MHD motif of Pikp-2 abolished cell death responses to AVR-PikD (Zdrzalek et al., 2020). Consequently, we introduced the equivalent mutation in Pikm-2 and verified that it also abrogated cell death in autoimmune combinations (**Figure 4**), confirming that Pikm-2 requires an intact MHD motif to trigger cell death and strengthening the link between constitutive cell death and immune activation.

### NLR specialization and autoimmunity are linked to the same amino acid polymorphism

Interestingly, only mismatches involving Pikm-2 triggered cell death in the absence of the effector (**Figure 3**), suggesting that this NLR harbours the determinants of this autoactive phenotype. To understand the basis of Pikm-2–mediated autoimmunity, we used the point mutants in Pik-2 polymorphic positions presented above **(Figure S3)** to explore the determinant of constitutive cell death. To this end, we co-expressed each mutant with either Pikp-1 or Pikm-1 in the presence or absence of AVR-PikD effector **(Figure 5, Figure S8)**. In this assay, we added AVR-PikD effectors as a positive control for cell death.

**Figure 5.**
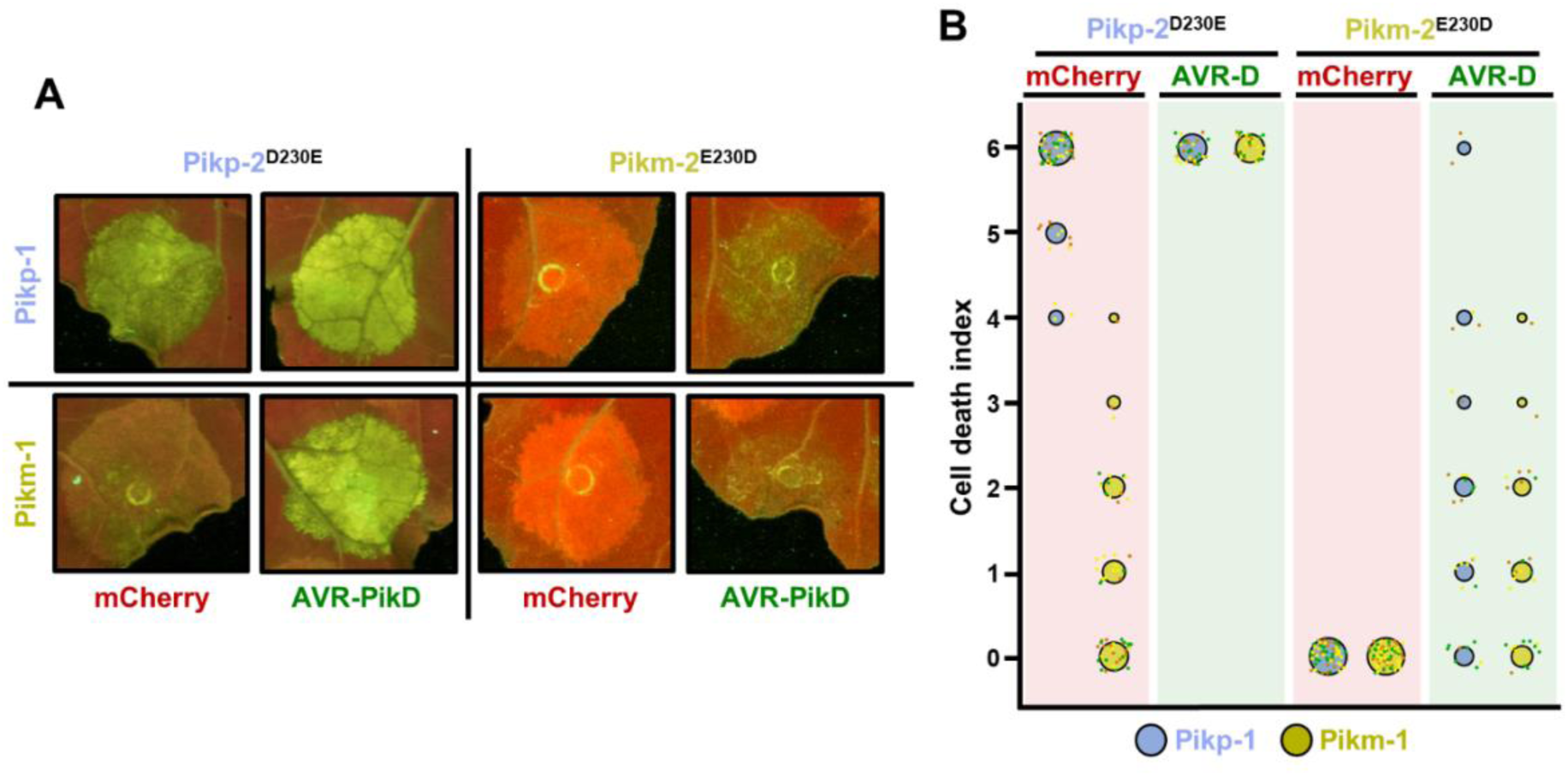
Polymorphism at position 230 in the NB-ARC domain is a Pik-2 determinant for constitutive cell death. **(A)** Representative leaf spot images and scoring of cell death mediated by Pik-2 as autofluorescence under UV-light. **(B)** Cell death scoring is represented as dot plots comparing cell death triggered by Pik-2 mutants at polymorphic positions 230. Pik-2 mutants were co-expressed with Pikp-1 (blue dots) or Pikm-1 (yellow dots) together with mCherry (red panel) or AVR-PikD (green panel). The number of repeats was 60 and 30 for the spots co-infiltrated with mCherry and AVR-PikD, respectively. For each sample, all the data points are represented as dots with a distinct colour for each of the three biological replicates; these dots are jittered about the cell death score for visualisation purposes. The size of the central dot at each cell death value is proportional to the number of replicates of the sample with that score.

The Asp230Glu mutation in Pikp-2 conferred a strong cell death response in the absence of the effector when co-expressed with Pikp-1, while only residual constitutive activation could also be observed with Pikm-1 (**Figure 5**). By contrast, the reciprocal mutation at the equivalent position in Pikm-2 abrogated constitutive cell death in the presence of Pikp-1 and reduced the cell death response mediated by AVR-PikD recognition (**Figure 5**). Single mutations in any of the other polymorphic positions had no effect on constitutive cell death activation **(Figure S8)**.

Additionally, we confirmed that constitutive cell death triggered by Pik-2 Asp230Glu is also dependent on the P-loop and MHD motifs, confirming that this mutation leads to immune activation **(Figure S9)**. Interestingly, cell death responses were reduced but not completely abolished when Pikm-2 or Pikp-2 Asp230Glu were co-expressed with a P-loop mutant of Pikp-1 **(Figure S10)**. Protein accumulation of the Pikp-1 P-loop mutant and the Pik-2 P-loop and MHD mutants were equivalent to their wild-type counterparts **(Figure S11)**. This unequal contribution of the P-loop motifs of sensor and helper NLRs adds an extra layer of information to the cooperation model of NLR activation previously proposed for Pik (Bialas et al., 2018).

Overall, we narrowed down a determinant of autoimmunity in the mismatched Pik pairs to a single amino acid polymorphism. Furthermore, we confirmed that this polymorphism mediates cell death phenotypes by a mechanism dependent on the P-loop and MHD motifs. Interestingly, the same polymorphism is related to the stronger cell death responses to AVR-Pik effectors mediated by Pikm (**Figure 2**). Altogether, this establishes a link between immune specialization and gain of constitutive cell death responses in NLR pairs, two hallmarks of coevolution.

### The Glu230 amino acid polymorphism has evolved in modern rice

Having identified a determinant of Pik NLR pair specialization and compatibility as a single amino acid polymorphism, we aimed to gain an evolutionary perspective of the specialization process of Pik-2. For this, we combined the Pik-2 coding sequences from rice cultivars described above with the Pik-2 orthologs from wild Asian and African relative species (Bialas et al., 2021; Stein et al., 2018) (see methods for accession numbers) and calculated the maximum likelihood phylogenetic tree rooted in the African outgroup species *Leersia perrieri* (**Figure 6A**).

**Figure 6.**
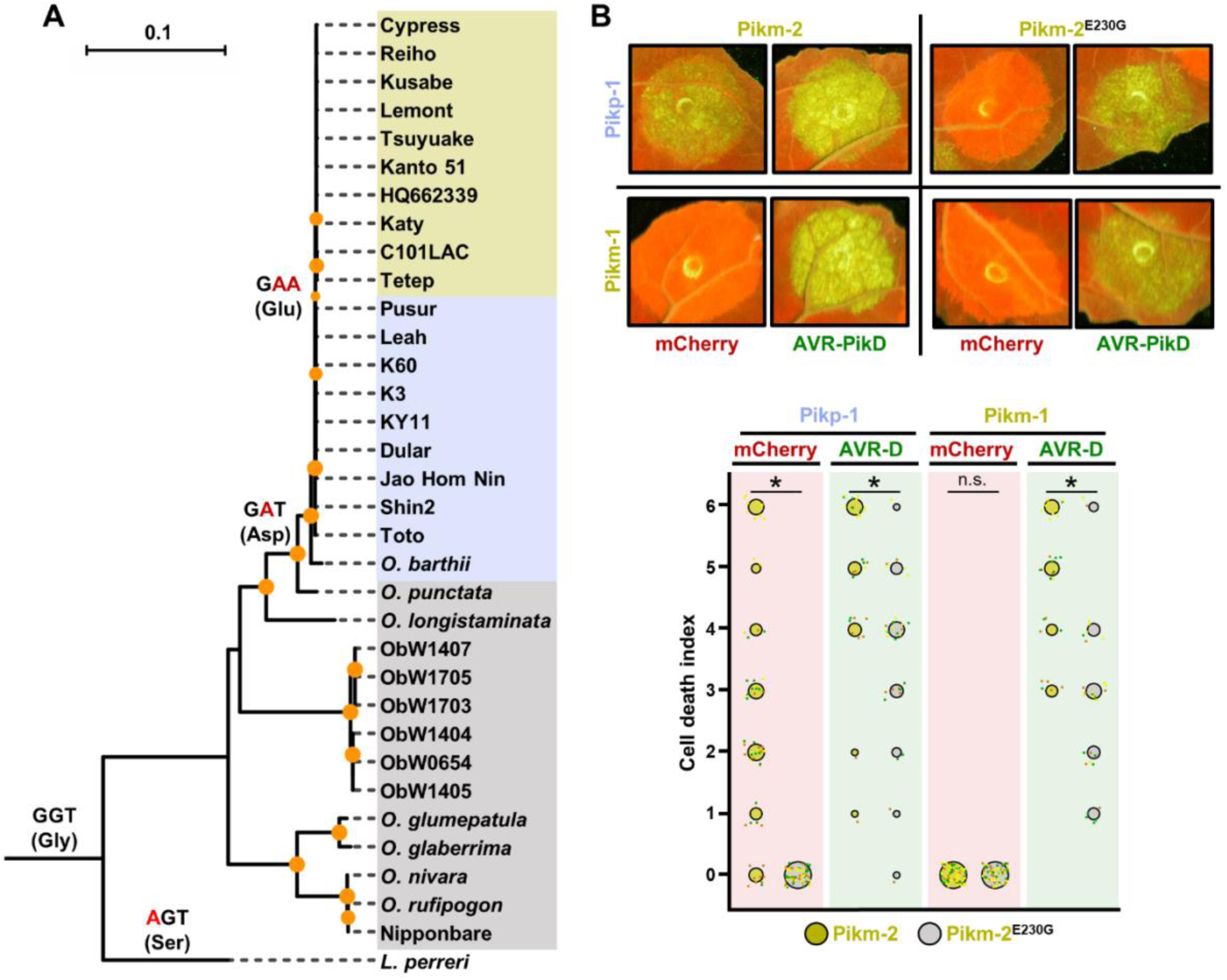
(A) Reconstruction of the evolutionary history of Pik-2 polymorphism at position 230. Maximum likelihood (ML) phylogenetic tree of Pik-2 coding sequences from cultivated rice and wild rice species. The tree was calculated from a 3,066-nt-long alignment using RAxML v8.2.11 (Stamatakis, 2014), 1000 bootstrap method (Felsenstein, 1985) and GTRGAMMA substitution model (Tavaré, 1986). Best-scoring ML tree was manually rooted using the Pik-2 sequence from *Leersia perreri* as an outgroup. The bootstrap values above 80 are indicated with orange circles at the base of respective clades; the support values for the relevant nodes are depicted by the size of the circle. The scale bar indicates the evolutionary distance based on the nucleotide substitution rate. The tree was represented using Interactive Tree Of Life (iTOL) v4 (Letunic and Bork, 2019). The tree shows a set of inferred nucleotides (states) at the Pik-2 polymorphic position 230 based on their predicted likelihood at sites 709 to 711 of the sequence alignment. Non-synonymous changes at the codon are depicted in red next to their corresponding node. For visualization, rice species and cultivars names are shaded in gold, light blue or grey according to their residue in Pik-2 polymorphic position 230 (Glu, Asp or Gly, respectively). Ob: *Oryza brachyantha*. **(B) Reversion to ancestral state of Pikm-2 Glu230 abolish autoimmunity.** Representative leaf spot images depicting Pik-mediated cell death as autofluorescence under UV-light in the presence or absence of AVR-Pik effector. Scoring of the cell death triggered by Pikm-2 or Pikm-2 Glu230Gly mutant when co-expressed with Pikp-1 or Pikm-1 is represented as dot plots. The number of repeats was 60 and 30 for the spots co-infiltrated with mCherry and AVR-PikD, respectively. For each sample, all the data points are represented as dots with a distinct colour for each of the three biological replicates; these dots are jittered about the cell death score for visualisation purposes. The size of the central dot at each cell death value is proportional to the number of replicates of the sample with that score. Significant differences between relevant conditions are marked with an asterisk and the details of the statistical analysis are summarised in **Figure S12**.

Pik-2 sequences from wild rice species are phylogenetically distinct from those belonging to modern rice, with the exception of Nipponbare and Toto (**Figure 6A**). These modern varieties make two distinct groups harbouring Pikp cultivar K60 or Pikm cultivar Tsuyuake (**Figure 6A**) (Bialas et al., 2021; De la Concepcion et al., 2021).

To learn more of the evolutionary trajectory of Pik-2, we inferred the ancestral state of the nucleotide sequences coding for the polymorphic position 230. This analysis revealed that a Gly residue encoded by GGT is an ancestral state at this position and is still present in most Pik-2 sequences from wild *Oryza* species (**Figure 6A**).

A transition from GGT (coding for Gly) to GAT (coding for Asp) in position 230 occurred before the split of *Oryza sativa* and *Oryza punctata* and has been maintained in Pik-2 NLRs of modern rice varieties clustering in the with the Pikp cultivar K60 (**Figure 6A**). This change opened the possibility of a non-synonymous Asp to Glu mutation by a GAT to GAA transversion, which occurred in the rise of the clade containing the Pikm cultivar Tsuyuake. This Asp230Glu polymorphism represents a specialization determinant in the Pikm NLR pair and ultimately rendered Pikm-2 incompatible with Pikp-1.

To experimentally validate the reconstructed evolutionary history of Pik-2 polymorphic position 230, we reverted this position in Pikm-2 to the ancestral state by introducing a Glu230Gly mutation and tested its ability to trigger cell death in *N. benthamiana*. The Glu230Gly mutation abolished the constitutive cell death triggered by Pikm-2 when co-expressed with Pikp-1 in the absence of the effector (**Figure 6B**). This mutation did not abrogate the cell death response to the AVR-PikD effector, although it slightly reduced it compared with the wild type **(Figure 6B, Figure S12)**. Protein accumulation of Pikm-2 Glu230Gly was equivalent to wild-type Pikp-2 and Pikm-2 **(Figure S13)**.

Overall, having reconstructed the evolutionary history of Pik NLR specialization we propose a model where a multi-step mutation led to the emergence of Glu230 polymorphism, which is linked to an efficient cell death response to AVR-Pik effectors in the Pikm pair. We further demonstrated that the rise of this polymorphism is associated with NLR incompatibility with mismatched sensor NLRs from the Pikp-like clade, triggering constitutive immune activation and cell death in the absence of pathogen effectors.

### Sensor/helper hetero-pairing alters protein accumulation in Pik NLRs

We aimed to obtain mechanistic understanding of Pik NLR pair coevolution and autoactivation. For this, we investigated whether accumulation of sensor Pik-1 or helper Pik-2 proteins is altered in the presence of the coevolved or mismatched pair. After co-expression of both Pikp-1 and Pikm-1 alleles in *N. benthamiana* in combination with the helper Pikp-2 or Pikm-2 alleles followed by western blot, we observed that protein accumulation of Pik-1 and Pik-2 alleles were consistently increased when they were expressed together compared to co-expression with empty vector (**Figure 7A**). This is consistent with a model where Pik-1 and Pik-2 associate in sensor/helper NLR heterocomplexes, stabilizing both proteins (Zdrzalek et al., 2020).

**Figure 7.**
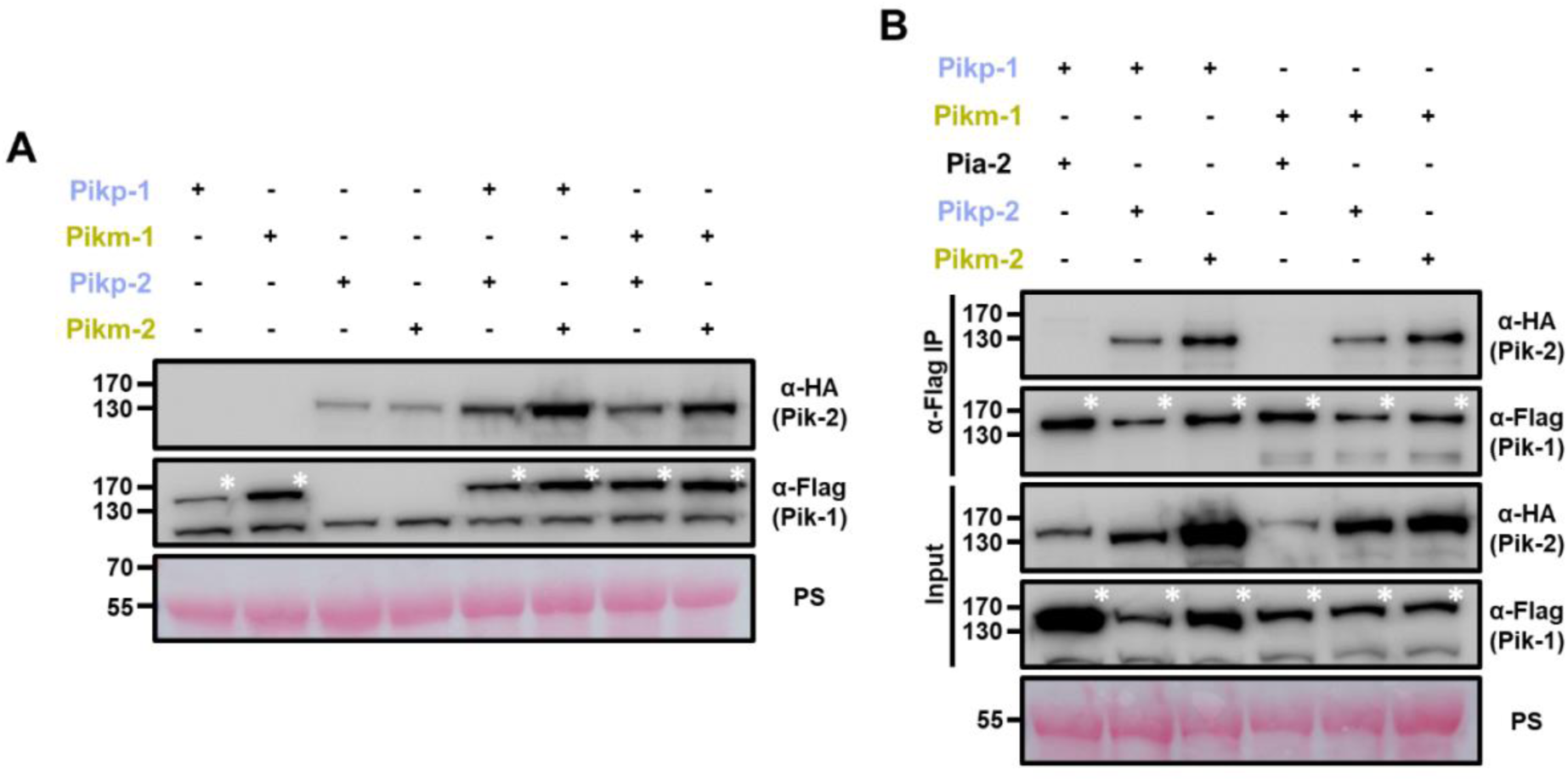
(A) Increased protein accumulation of paired Pik proteins when co-expressed together in planta. Western blots showing protein accumulation of Pik-1 and Pik-2 alleles in different combinations. C-terminally 6*×*His3*×*FLAG tagged Pik-1 alleles were transiently co-expressed with empty vector (EV) or C-terminally 6*×*HA tagged Pik-2 alleles in *N. benthamiana*. Total protein extracts were probed with α-FLAG and α-HA antisera for Pik-1 and Pik-2, respectively. Asterisks mark the band corresponding to Pik-1. **(B) Mismatched Pik NLR pairs associate in planta.** Co-immunoprecipitation of full-length Pikp-1 and Pikm-1 alleles in combination with either Pikp-2 or Pikm-2 helper NLRs. C-terminally 6*×*HA tagged Pia-2, Pikp-2 or Pikm-2 NLRs were transiently co-expressed with Pikp-1:6*×*His3*×*FLAG or Pikm-1:6*×*His3*×*FLAG in *N. benthamiana*. Immunoprecipitates obtained with anti-FLAG antiserum, and total protein extracts, were probed with appropriate antisera. Asterisks mark the band corresponding to Pik-1. Total protein loading is shown by Ponceau staining (PS).

Interestingly, accumulation of the helper Pik-2 in the autoimmune pair Pikp-1/Pikm-2 was consistently higher (**Figure 7A**). This could be due to a different sensor/helper stoichiometry in the constitutively active Pik complex, as observed in some activated NLR complexes (Hu et al., 2015; Sharif et al., 2019; Tenthorey et al., 2017; Zhang et al., 2015). This is also consistent with the finding that CC domain of Pik-2 NLR has the consensus MADA motif first identified in ZAR1 (Adachi et al., 2019a), indicating the possibility that Pik activation may involve oligomerization of multiple Pik-2 receptors as in the ZAR1 resistosome (Wang et al., 2019a).

### Coevolved and mismatched Pik pairs form heterocomplexes

Prompted by the differences in protein accumulation observed between different combinations of Pik-1 and Pik-2, we investigated whether cell death phenotypes in mismatched Pik pairs are underpinned by differences in NLR hetero-association.

We co-expressed C-terminally tagged Pikp-1 or Pikm-1 with either C-terminally tagged Pikp-2 or Pikm-2 in *N. benthamiana*. Following total protein extraction, we performed co-immunoprecipitation to test for differences in NLR association (**Figure 7B**). Pikp-1 and Pikm-1 were also co-infiltrated with the rice NLR Pia-2 (the sensor NLR, also known as RGA5, of the immune receptor pair Pia) as a negative control.

Both Pikp-2 and Pikm-2 could be detected after immunoprecipitation of either Pikp-1 or Pikm-1 sensor NLRs (**Figure 7B**). Additionally, none of the Pik-2 mutations generated above seem to have a measurable effect on the sensor/helper association **(Figure S14, S15)**.

These results indicate that cell death phenotypes observed in mismatched pairs are not underpinned by major alterations in association. Instead, Pik sensor and helper NLRs may form pre-activation complexes in the resting state and subtle changes, perhaps both in association and stoichiometry between Pik NLRs, govern cell death responses and autoimmune phenotypes described above.

### Sensor/helper association of Pik NLR pairs is independent of NLR activation

As Pik NLR pairs associate in pre-activation complexes (**Figure 7B**) (Zdrzalek et al., 2020), we investigated whether this process requires functional NLRs. We co-expressed the Pikm-2 P-loop and MHD mutants with either Pikp-1 or Pikm-1 in *N. benthamiana*. Following protein extraction and immunoprecipitation of Pik-1, we found that these mutations do not affect the ability to associate with the sensor NLR Pik-1 compared to wild-type Pikm-2 (**Figure 8**), although they completely abolish Pik-mediated cell death. Similarly, the reduced cell death activity in the Pik-1 P-loop mutant did not correlate with alterations in the association to the helper NLR Pik-2 (**Figure S16**).

**Figure 8.**
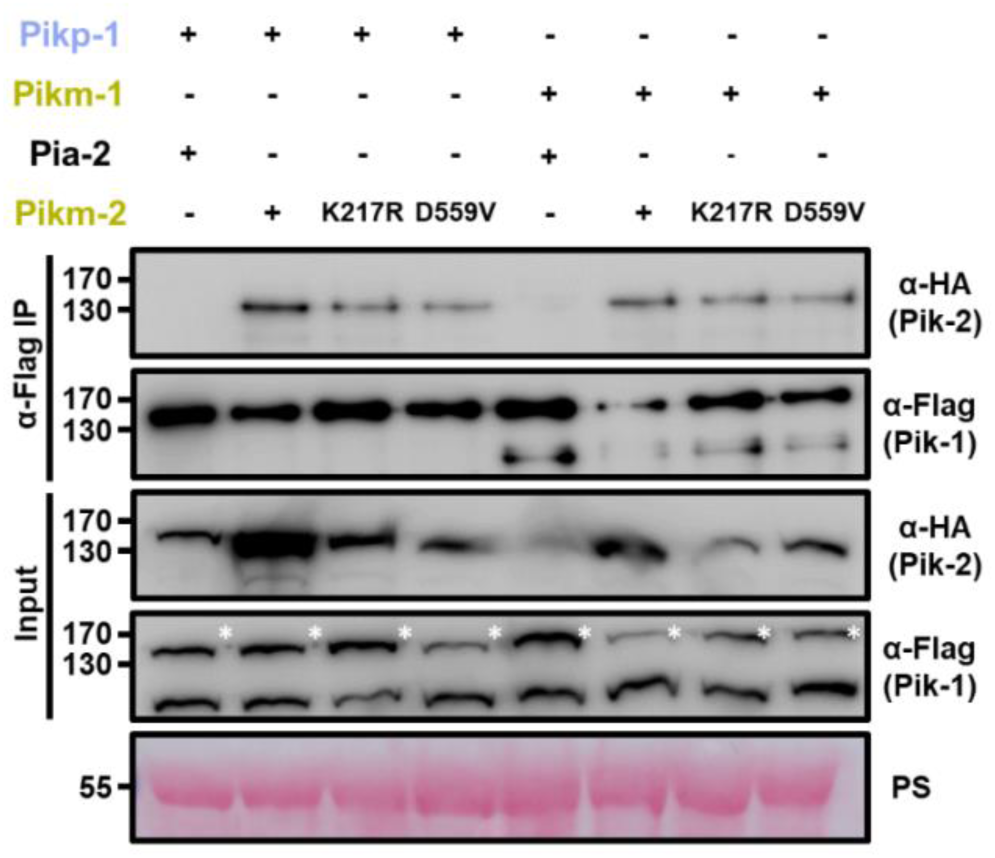
Mutations in Pik-2 P-loop and MHD motifs do not affect in planta association of Pik-1. Co-immunoprecipitation of Pikm-2 P-loop and MHD mutants with full length Pikp-1 and Pikm-1 alleles. C-terminally 6*×*HA tagged Pikm-2 mutants in P-loop (Lys217Arg) and MHD (Asp559Val) motifs were transiently co-expressed with either Pikp-1:6*×*His3*×*FLAG or Pikm-1:6*×*His3*×*FLAG in *N. benthamiana*. Immunoprecipitates obtained with anti-FLAG antiserum, and total protein extracts, were probed with appropriate antisera. Co-expression with C-terminally tagged 6*×*HA Pia-2 NLR and wild-type Pikm-2 were included as negative and positive control, respectively. Asterisks mark the band corresponding to Pik-1. Total protein loading is shown by Ponceau staining (PS).

These results imply that pre-activated Pik NLR pair association does not require functional NLRs and is independent of nucleotide binding. In the native state such pre-activation complexes may require ADP/ATP exchange to induce or stabilise changes in receptor conformation and/or stoichiometry to trigger immune signalling.

### Sensor and helper Pik NLRs preferentially associate with their coevolved pair

To gain a deeper knowledge of Pik pair association, we investigated whether allelic Pik NLRs display any preference in association to their coevolved NLR pair. As both autoactive and non-autoactive pairs associate, we designed an NLR competition assay with a cell death readout to test for preferential association between allelic NLRs **(Figure S17)**. For this, we took advantage of the constitutive cell death phenotype triggered by the association of Pikp-1 and Pikm-2 **(Figure S17A)**. In a scenario where a non-autoactive Pik-2 NLR displays higher helper/sensor association to Pikp-1, Pikm-2 would be outcompeted from complex formation, reducing the levels of constitutive cell death **(Figure S17B)**.

To test this, we transiently co-expressed both Pikp-1 and Pikm-2 NLRs in *N. benthamiana* using a fixed concentration (OD_600_ 0.4) of *Agrobacterium tumefaciens* to deliver each construct. We also co-delivered increasing concentrations of Pikp-2 (spanning an OD_600_ of 0–0.6) and scored the cell death phenotype (**Figure 9**).

**Figure 9.**
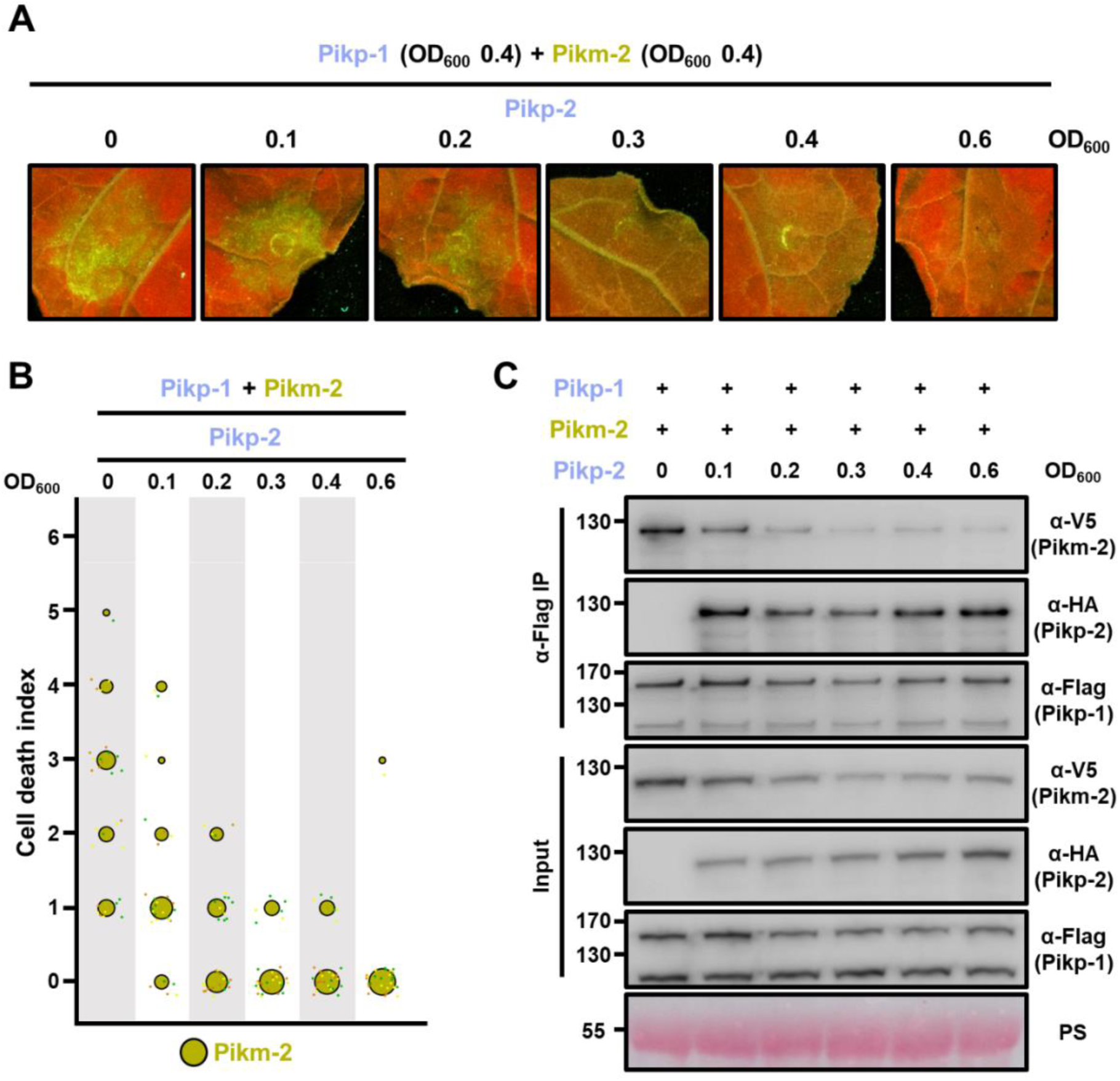
Pikp-2 supresses constitutive cell death mediated by Pikm-2. **(A)** Representative leaf spot images depicting Pikm-2 mediated cell death in the presence of Pikp-1 and increasing concentration of Pikp-2 as autofluorescence under UV-light. For each experiment, Pikp-1 and Pikm-2 were co-infiltrated at OD_600_ 0.4 each. Increasing concentrations of Pikp-2 were added to each experiment (from left to right: OD_600_ 0, 0.1, 0.2, 0.3, 0.4 and 0.6). **(B)** Scoring of the cell death assay is represented as dot plots. A total of three biological replicates with 10 internal repeats each were performed for each experiment. For each sample, all the data points are represented as dots with a distinct colour for each of the two biological replicates; these dots are jittered about the cell death score for visualisation purposes. The size of the central dot at each cell death value is proportional to the number of replicates of the sample with that score. **(C) Pikp-2 outcompetes Pikm-2 association to Pikp-1.** Co-immunoprecipitation of Pikm-2 and Pikp-1 in the presence of increasing concentrations on Pikp-2. C-terminally V5 tagged Pikm-2 and C-terminally 6*×*His3*×*FLAG tagged Pikp-1 were transiently co-expressed in *N. benthamiana* alongside with increasing concentrations of C-terminally 6*×*HA tagged Pikp-2 (from left to right: 0, 0.1, 0.2, 0.3, 0.4 and 0.6 OD_600_). Immunoprecipitates obtained with anti-FLAG antiserum, and total protein extracts, were probed with appropriate antisera. Asterisks mark the band corresponding to Pikp-1. Total protein loading is shown by Ponceau staining (PS).

Interestingly, Pikp-2 acted as a suppressor of autoimmune phenotypes triggered by Pikp-1/Pikm-2 as increasing concentrations of Pikp-2 lowered the constitutive cell death phenotype (**Figure 9A, B**). This reduction in cell death was evident even in the lowest concentration of Pikp-2 (**Figure 9A, B**), suggesting that Pikp-1 displays preference to signal through coevolved Pikp-2 rather than Pikm-2.

We also replicated this experiment co-infiltrating a fixed concentration of Pikp-1 and Pikp-2, with increasing concentration of Pikm-2. In agreement with a signalling preference between Pikp-1 and Pikp-2, Pikm-2 could not overcome the suppression by the presence of Pikp-2, even at the highest concentration **(Figure S18)**.

To investigate whether the decrease in cell death is correlating with reduced association of the maladapted pair Pikp-1/Pikm-2 in the presence of Pikp-2, we immunoprecipitated Pikp-1 and tested for the presence of Pikp-2 or Pikm-2 (**Figure 9C**).

Differences in protein accumulation observed in the different sensor/helper combinations of Pik pairs makes it particularly challenging to obtain even inputs for this experiment. As reported above (**Figure 7A**), the Pik-2 proteins are more stable in association with Pik-1, therefore, if a Pik-2 protein is outcompeted from a hypothetical complex, it will present reduced accumulation in the input. The contrary effect occurs in the Pik-2 proteins forming autoactive complexes, as they showed increased accumulation in autoactive combinations (**Figure 7A**), the amount of protein in concentrations where Pikp-2 supresses constitutive cell death may seem lowered.

Nevertheless, co-immunoprecipitation results depicted a preference in association of Pikp-1 to Pikp-2 over Pikm-2. Increasing concentrations of Pikp-2 reduced the association of Pikp-1 to Pikm-2, outcompeting Pikm-2 from a heterocomplex with Pikp-1 (**Figure 9C**). This correlates with the reduction of the constitutive cell death assay observed in the NLR competition experiments (**Figure 9A, B**).

Altogether, these data reveal that coevolved Pik NLRs display preference in association over non–coevolved NLRs. This represents another example of NLR pair co-adaptation. These differences may underpin the observed cell death phenotypes in response to effectors and in autoimmunity.

### Pik helper/sensor association preference is underpinned by Pik-2 polymorphism

To shed light on the basis of the preferential binding between Pikp-1 and Pikp-2, we tested the role of the polymorphism 230 in this phenotype. For this, we repeated the NLR competition assay co-infiltrating a fixed concentration (OD_600_ 0.4) of Pikp-1 and the autoactive mutant Pikp-2 Asp230Glu, with increasing amounts of Pikp-2.

The combination of Pikp-1 and Pikp-2 Asp230Glu led to a strong cell death in the absence of effector (**Figure 5**). However, increasing concentrations of Pikp-2 significantly reduced this phenotype (**Figure 10**).

**Figure 10.**
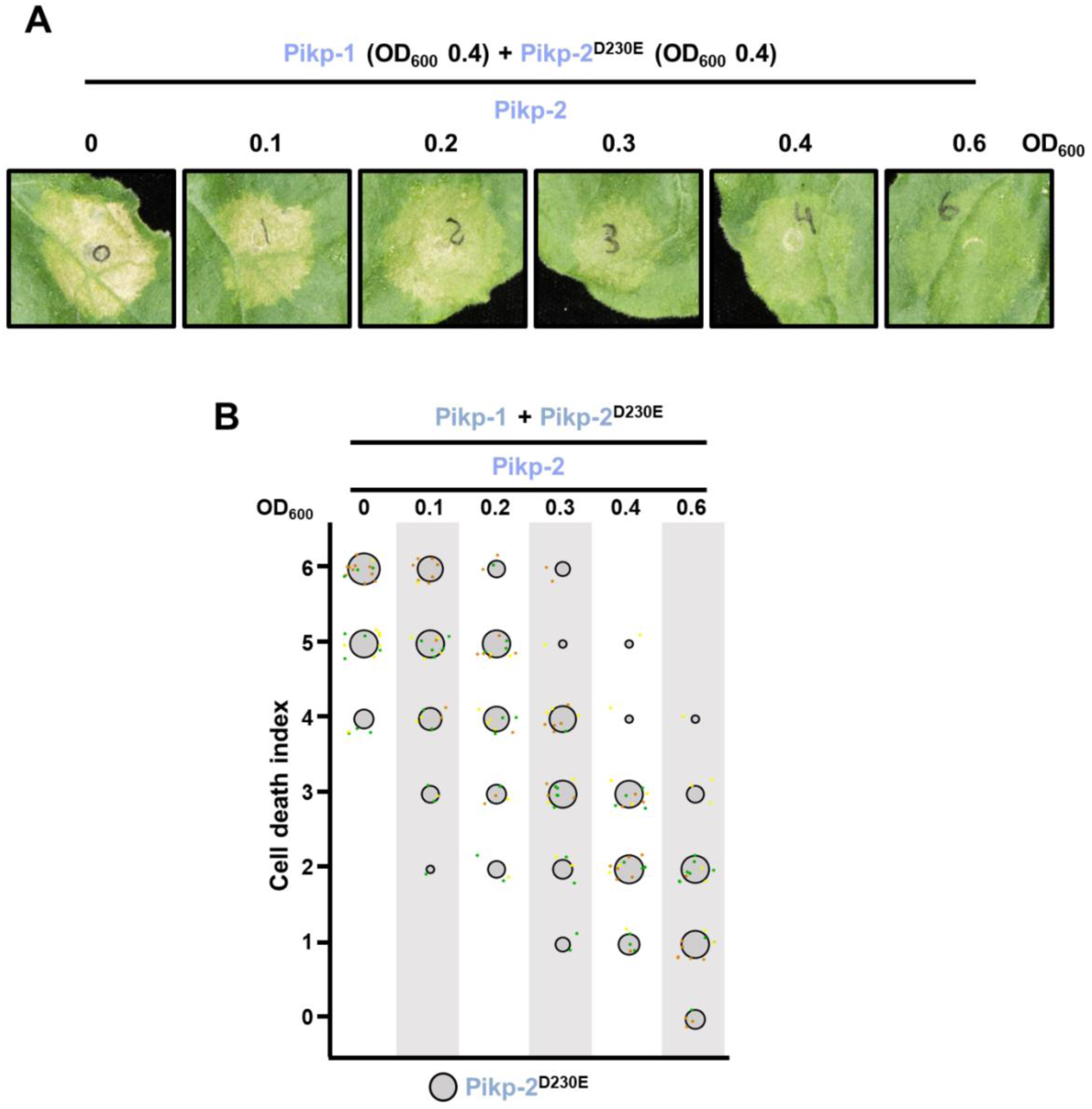
Wild-type Pikp-2 supresses constitutive cell death mediated by Pikp-2 Asp230Glu mutant. **(A)** Representative leaf spot images depicting Pikp-2 Asp230Glu mediated cell death in the presence of Pikp-1 and increasing concentration of Pikp-2. For each experiment, Pikp-1 and Pikp-2 Asp230Glu were co-infiltrated at OD_600_ 0.4 each. Increasing concentrations of Pikp-2 were added to each experiment (from left to right: OD_600_ 0, 0.1, 0.2, 0.3, 0.4 and 0.6). **(B)** Scoring of the cell death mediated by Pikp-2 Asp230Glu in the presence of Pikp-1 and increasing concentration of Pikp-2 assay represented as dot plots. For each experiment, Pikp-1 and Pikp-2 Asp230Glu were co-infiltrated at OD_600_ 0.4 each. Increased concentration of Pikp-2 was added to each experiment (from left to right: OD_600_ 0, 0.1, 0.2, 0.3, 0.4 and 0.6). A total of three biological replicates with 10 internal repeats each were performed for each experiment. For each sample, all the data points are represented as dots with a distinct colour for each of the three biological replicates; these dots are jittered about the cell death score for visualisation purposes. The size of the central dot at each cell death value is proportional to the number of replicates of the sample with that score.

This indicates the Pik-2 Glu230 polymorphism may also be related to the preferential association between sensor and helper NLRs in addition to its role in specialization towards AVR-Pik effector response and autoimmunity.

### Pik pair preferential association requires NLR activation

To investigate if the preferential sensor/helper association is related to the activation of the helper NLR Pik-2, we tested whether the constitutive cell death mediated by Pikp-2 Asp230Glu could be supressed by mutants that render Pikp-2 inactive.

Although Pik NLRs do not require a functional P-loop or MHD motif to form heterocomplexes (**Figure 8**), we did not observe reduction of cell death phenotypes with increasing concentrations of Pikp-2 mutants in the P-loop (Lys217Arg) or MHD (Asp559Val) motifs (**Figure 11**), even at the highest concentration. This indicates that the P-loop and MHD motifs are required for the preferential sensor/helper association observed in Pikp-1 and Pikp-2.

**Figure 11.**
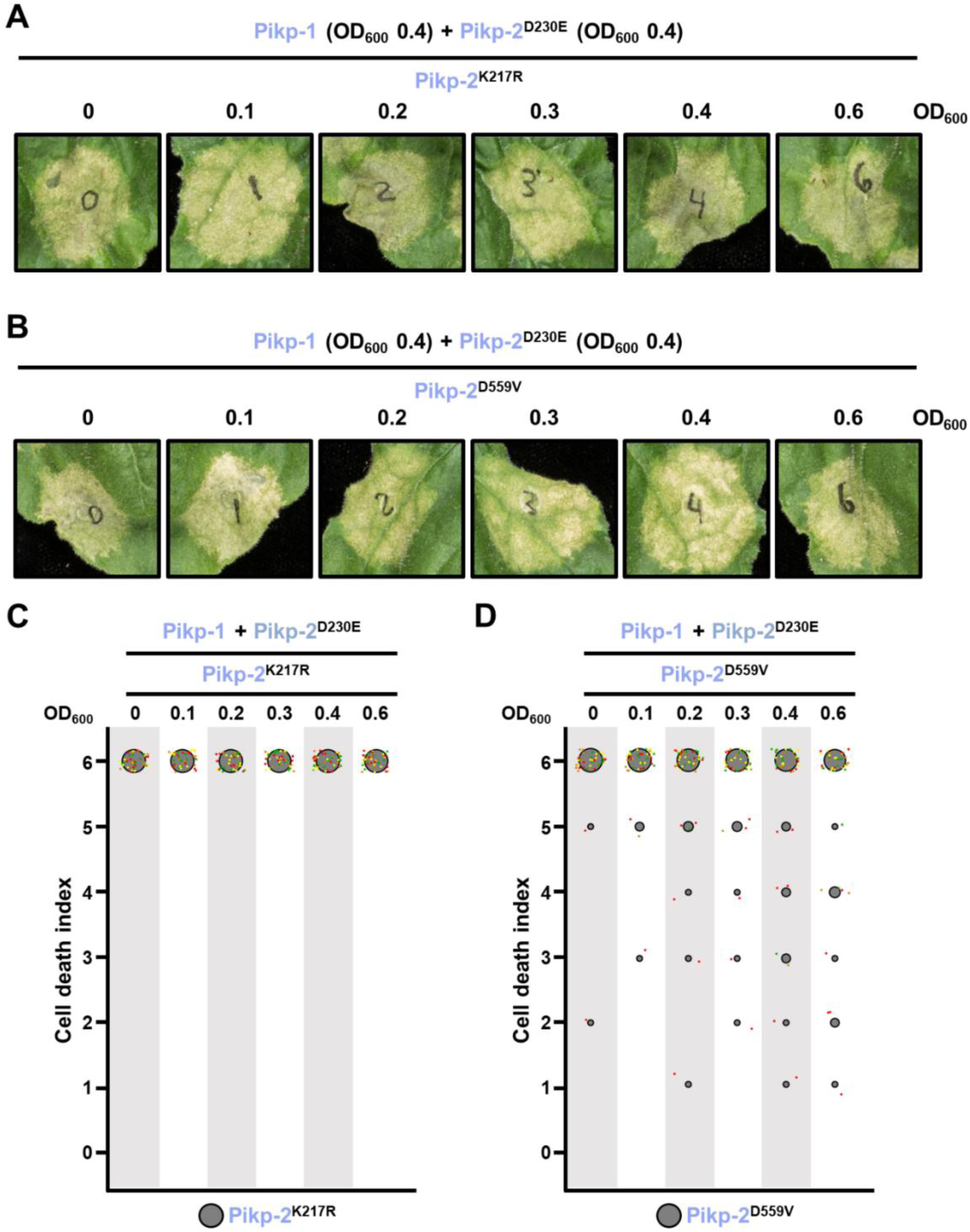
Suppression of constitutive cell death mediated by Pikp-2 Asp230Glu requires an active Pikp-2. Representative leaf spot images depicting Pikp-2 Asp230Glu mediated cell death in the presence of Pikp-1 and increasing concentration of Pikp-2. For each experiment, Pikp-1 and Pikp-2 Asp230Glu were co-infiltrated at OD_600_ 0.4 each. Increasing concentrations of **(A)** Pikp-2 Lys217Arg or **(B)** Pikp-2 Asp559Val were added to each experiment (from left to right: OD_600_ 0, 0.1, 0.2, 0.3, 0.4 and 0.6). Scoring of the cell death mediated by Pikp-2 Asp230Glu in the presence of Pikp-1 and increasing concentration of **(C)** Pikp-2 Lys217Arg or **(D)** Pikp-2 Asp559Val represented as dot plots. For each experiment, Pikp-1 and Pikp-2 Asp230Glu were co-infiltrated at OD_600_ 0.4 each. Increased concentration of Pikp-2 mutants were added to each experiment (from left to right: OD_600_ 0, 0.1, 0.2, 0.3, 0.4 and 0.6). A total of four biological replicates with 10 internal repeats each were performed for each experiment. For each sample, all the data points are represented as dots with a distinct colour for each of the four biological replicates; these dots are jittered about the cell death score for visualisation purposes. The size of the central dot at each cell death value is proportional to the number of replicates of the sample with that score.

Altogether, these results suggest that changes in Pik sensor and helper NLR association from a resting state into an activated complex requires functional NLRs. This is consistent with studies in the Arabidopsis NLR RPP7, where a P-loop mutant retains the ability to associate with autoactive forms of its incompatibility partner HR4 but is not capable of forming higher order assemblies (Li et al., 2020).

## Discussion

The work presented here highlights sensor/helper coevolution in an allelic rice NLR pair and the basis of their functional diversification towards differential effector recognition specificities (**Figure 12**). We discovered that a single amino acid polymorphism underpins specialization of the helper Pik-2 NLR to its corresponding Pik-1 sensor NLR. Changes in this residue affect cell death outcomes in effector recognition and autoimmune phenotypes. By narrowing down the contribution of NLR specialization to a single amino acid, we could trace the evolutionary history of this polymorphism (**Figure 12**).

**Figure 12.**
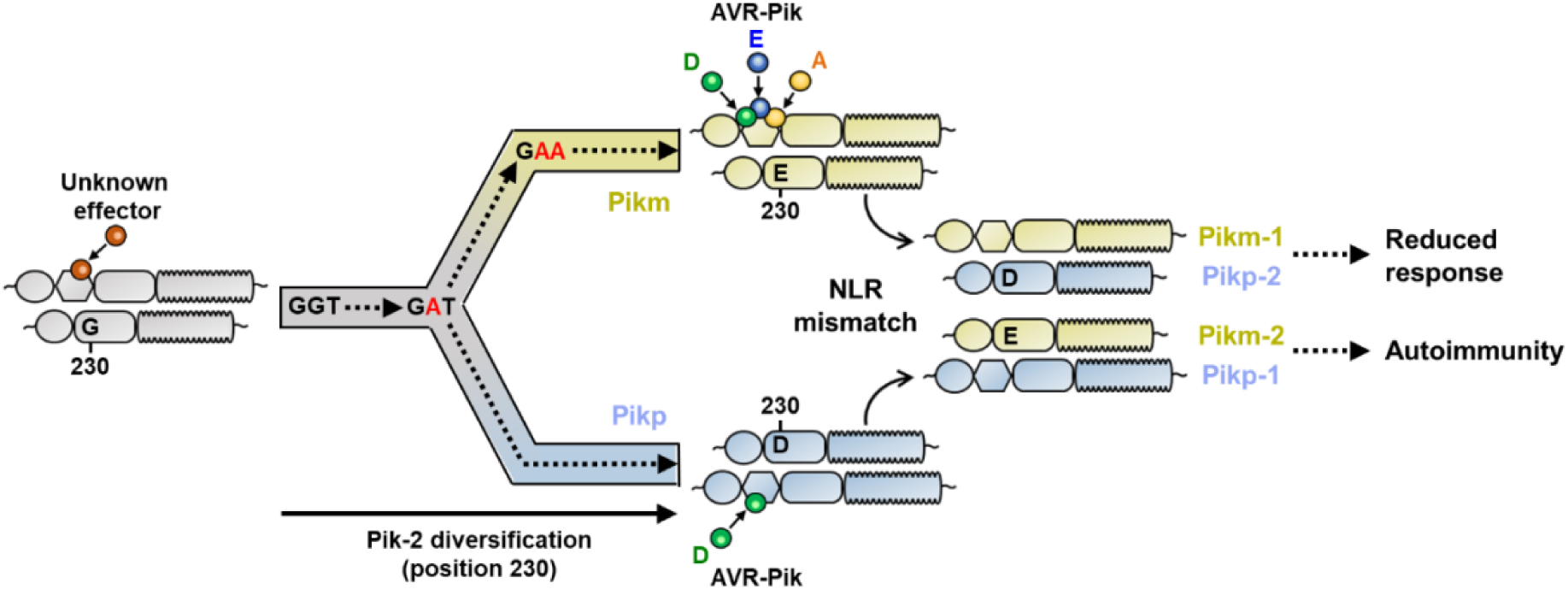
Schematic representation of the proposed evolutionary model of the Pik pairing. Pikp (coloured in ice blue) and Pikm (coloured in gold), have evolved and specialized from an ancestral NLR pair (coloured in grey), functionally diversifying and gaining recognition to a different subset of allelic AVR-Pik effectors. Residues at Pik-2 polymorphic position 230 are indicated and mutations predicted to have occurred during this transition are indicated in red. As a consequence of diversification, mismatch of Pikp and Pikm impairs immune responses and leads to NLR autoactivation and constitutive cell death in *N. benthamiana*.

The notion that NLRs can work together in pairs is now well-established in the field of plant–microbe interactions (Adachi et al., 2019b; Jubic et al., 2019). Under this emerging framework, it is predicted that cooperating NLRs co-adapt to optimise and maintain a tight control over immune responses. However, the extent to which paired NLRs coevolve to efficiently respond to pathogen effectors while keeping a fine-tuned regulation of immune responses is not well understood at the molecular level. Particularly intriguing is how rapid changes driven by coevolution with pathogen effectors and major evolutionary events, such as the integration of an unconventional domain, impact NLR co-adaptation.

The genetic linkage of the Pik NLR pair has been maintained in grass genomes for tens of millions of years and emerged before the integration of the HMA domain in Pik-1 (Bialas et al., 2021). This suggests that Pik-1 and Pik-2 have been coevolving for a long time, potentially before providing resistance to the blast fungus. Thus, the integration of the HMA domain in Pik-1, and its subsequent rapid coevolution with rice blast effectors, may have represented a major perturbation on the coevolutionary equilibrium in the paired Pik NLRs. Here we demonstrated how allelic Pik NLR pairs have differentially coevolved and functionally specialized, leading to autoimmune phenotypes when mismatched. This suggests that in response to HMA integration and diversification in the sensor NLR Pik-1, its helper NLR Pik-2 has acquired polymorphisms to avoid loss of function and/or triggering autoimmunity.

To date, integrated domains have been primarily found in paired NLRs that are located in co-regulatory modules with a shared promoter region (Cesari et al., 2014a). Therefore, the spatial regulation of NLRs with unconventional domains in pairs might be a general mechanism to mitigate NLR misregulation as a consequence of domain integrations or their accelerated evolutionary rates compared with other NLR domains (Bialas et al., 2021).

We found mismatched allelic NLR pairs can lead to constitutive cell death. We further narrowed down this phenotype to a single Asp to Glu polymorphism, which is the same polymorphism that underpins an extended cell death response to AVR-Pik effectors. Introducing this Asp230Glu polymorphism in Pikp-2 led to an increase of cell death in response to AVR-Pik effectors as well as to autoimmune phenotypes. As these amino acids have very similar properties it is intriguing how a fairly minor difference can underpin such a major phenotype. The mechanistic basis of this autoactivation phenotype remains obscure, but it is possible that the larger amino acid side chain (Glu carries an extra methylene group in the side chain) is sufficient to perturb protein–protein interactions that support transition to the active state of the NLR pair. Analogous Asp to Glu changes have been previously shown to act as a gain-of-function mutation in response regulators and transcription factors (Sakai et al., 2001; To et al., 2007). In some cases, the Asp residue is a target of phosphorylation and the change to Glu partially acts as a phosphomimetic mutation that leads to an active form (Klose et al., 1993). However, to date, there is no evidence to suggest that phosphorylation of Asp230 is involved in Pikp-2 activation. The Asp to Glu change did not prevent sensor/helper association, although it affected association preference. Altogether, this illustrates that small changes in NLR receptors can have profound phenotypical effects on immune regulation and cell death responses.

We still lack detailed information about the activation mechanism of paired plant NLRs although a cooperation mechanism has been proposed for the Pik NLR pair (Zdrzalek et al., 2020). By taking advantage of constitutively active Pik sensor/helper combinations and mutants we can expand our knowledge of NLR signalling mechanisms. The use of constitutively active immune receptors as a research tool is starting to be explored in the field of NLR biology. This approach has the advantage of simplifying the complex requirements of immune activation by removing the variability of the effector. It also renders full receptor activation, whilst relying solely on effector recognition can provide a mixture of active and inactive receptors.

Sensor and helper Pik NLRs form a pre-activation complex (Zdrzalek et al., 2020). Activation of immune responses may rearrange the composition of this complex, possibly affecting sensor/helper stoichiometry, as described for NAIP/NLRC4 inflammasomes (Hu et al., 2015; Tenthorey et al., 2017; Zhang et al., 2015). This rearrangement is dependent on nucleotide binding and has been fine-tuned during the evolutionary process, as depicted in the competition assays in the Pik pair.

Autoactive mutations in Pikp-2 led us to re-evaluate the involvement of conserved p-loop and MHD motif of sensor and helper Pik NLRs in signalling activation. In contrast to the previously described cooperation mechanism of Pik regulation (Bialas et al., 2018; Zdrzalek et al., 2020), the p-loop of sensor NLR Pik-1 is important but not necessary for NLR activation. This also deviates from the negative regulation mechanism described for other NLR pairs such as Pia or Arabidopsis RRS1/RPS4 (Cesari et al., 2014b; Cesari et al., 2013; Le Roux et al., 2015; Sarris et al., 2015) and suggests the Pik pair may trigger cell death via a different mechanism. Future experiments using autoactive combinations should help unravel the requirements of Pik NLR immune signalling and will reveal the nature of the changes undergone by NLRs during activation.

In summary, this work provides an evolutionary framework for how differential selective pressures, such as recognition of pathogen strains via effector binding, impact NLRs pairs. It uncovers the potential of paired NLRs to give rise to autoimmune phenotypes during evolution and links pathogen perception and autoimmunity.

## Materials and Methods

### Phylogenetics analyses

Codon-based alignment was generated using MUSCLE 3.8.425 (Edgar, 2004). The alignment positions with more than 40% data missing were removed using QKphylogeny (https://github.com/matthewmoscou/QKphylogeny). The maximum likelihood tree was calculated from a 3,066-nt-long alignment using 1000 bootstrap method (Felsenstein, 1985) and GTRGAMMA substitution model (Tavaré, 1986) as implemented in RAxML v8.2.11 (Stamatakis, 2014). Best-scoring tree was manually rooted using the Pik-2 sequence from *Leersia perreri* and visualized using the iToL tool v5.5.1 (Letunic and Bork, 2019). The interactive tree is publicly available at: https://itol.embl.de/tree/8229133147185181615486010

Joint reconstruction of ancestral sequences (Yang et al., 1995), based on the algorithm of Pupko et al. (Pupko et al., 2000), was performed using the codeml program as part of the PAML 4.9j package (Yang, 1997). The ancestral sequence reconstruction was carried out based on best-scoring ML tree and a 3,261-nt-long codon alignment of the full length Pik-2 sequences.

The accession numbers of the sequences used in the phylogenetic analyses are LPERR11G19580.2, ONIVA11G22700, ORUFI11G24740, XM_015762499.2, OGLUM11G22330, ORGLA11G0185700, MW568036, MW568041, MW568042, MW568043, MW568044, MW568045, KN541092.1, OPUNC11G19560, OBART11G23160.1, GU811862, HQ606329, HM048900_1, HQ662329_1, GU811867, AB462325, GU811861, GU811864, GU811865, GU811866, HQ662330, HM035360, KU365338.1, HQ660231, GU811868, GU811869, GU811870, GU811871, GU811872.

### Gene cloning

For protein expression in planta, we used full length Pikp-1 and Pikm-1 into the plasmid pICH47742 with a C-terminal 6*×*His/3*×*FLAG tag as previously described (De la Concepcion et al., 2018). Wild-type Pikp-2 and Pikm-2 in pICH47751 with C-terminal 6*×*HA were also described in (De la Concepcion et al., 2018) and Pik-2 mutated versions were generated by site-directed mutagenesis (see below) using appropriate Pik-2 template in pCR8/GW/TOPO (Invitrogen) with Golden Gate compatible overhangs. The constructs were later assembled in pICH47751 under control of *Agrobacterium tumefaciens* mannopine synthase (Mas) promoter and terminator and a C-terminal 6*×*HA using golden gate cloning (Engler et al., 2014). AVR-Pik effector alleles used in this study were previously described in (De la Concepcion et al., 2018).

All DNA constructs were verified by sequencing.

### Site-directed mutagenesis

Point mutations were introduced in Pik-2 by PCR amplification with Phusion™ polymerase (Thermo Fisher Scientific) using 5’-phosphorylated primers carrying the desired mutations. The amplification used primers running in opposite directions from the mutation site in the template Pik-2 in pCR8/GW/TOPO vector (Invitrogen). DNA templates were then eliminated by incubating the reaction with DpnI (New England Biolabs) for 1h at 37 °C. After PCR purification of the amplified products, the DNA sequence was re-ligated using T4 DNA ligase (New England Biolabs) according to the manufacturer’s protocol in 20 μl reactions incubated overnight at room temperature. Competent *E. coli* DH5α cells were subsequently transformed with 5 μl of the reaction. The resulting constructs were sequenced to ensure that a correct mutation was inserted into the sequence.

### in planta co-immunoprecipitation (Co-IP)

Transient gene-expression in planta for Co-IP was performed by delivering T-DNA constructs with *A. tumefaciens* GV3101 (C58 (rifR) Ti pMP90 (pTiC58DT-DNA) (gentR) Nopaline (pSoup-tetR)) strain into 4-week-old *N. benthamiana* plants grown at 22–25°C with high light intensity. *A. tumefaciens* strains carrying the given wild-type or mutated Pik-1 or Pik-2 were infiltrated at OD_600_ 0.2 each (unless otherwise stated), in agroinfiltration medium (10 mM MgCl_2_, 10 mM 2-(N-morpholine)-ethanesulfonic acid (MES), pH 5.6), supplemented with 150 µM acetosyringone.

For detection of complexes in planta, leaf tissue was collected 2–3 days post infiltration (dpi), frozen, and ground to fine powder in liquid nitrogen using a pestle and mortar. Leaf powder was mixed with 2 times weight/volume ice-cold extraction buffer (10% glycerol, 25 mM Tris pH 7.5, 1 mM EDTA, 150 mM NaCl, 2% w/v PVPP, 10 mM DTT, 1*×* protease inhibitor cocktail (Sigma), 0.1% Tween 20 (Sigma)), centrifuged at 4,200g at 4°C for 20–30 min, and the supernatant was passed through a 0.45 μm Minisart® syringe filter. The presence of each protein in the input was determined by SDS-PAGE/western blot. Wild-type and mutated Pik-1 and Pik-2 proteins were detected probing the membrane with anti-FLAG M2 antibody (Sigma) and anti-HA high affinity antibody 3F10 (Roche), respectively. For detection of Pikm-2 in the competition experiments in planta, we used anti-V5 antibody HRP-conjugated (Invitrogen).

For immunoprecipitation, 1.5 ml of filtered plant extract was incubated with 30 μl of M2 anti-FLAG resin (Sigma) in a rotatory mixer at 4°C. After three hours, the resin was pelleted (800g, 1 min) and the supernatant removed. The pellet was washed and resuspended in 1 ml of IP buffer (10% glycerol, 25 mM Tris pH 7.5, 1 mM EDTA, 150 mM NaCl, 0.1% Tween 20 (Sigma)) and pelleted again by centrifugation as before. Washing steps were repeated 5 times. Finally, 30 μl of LDS Runblue® sample buffer was added to the agarose and incubated for 10 min at 70°C. The resin was pelleted again, and the supernatant loaded on SDS-PAGE gels prior to western blotting. Membranes were probed with anti-FLAG M2 antibody (Sigma) and anti-HA high affinity antibody 3F10 (Roche) monoclonal antibodies. For competition experiments, the membrane was additionally probed with anti-V5 antibody HRP-conjugated (Invitrogen) to detect Pikm-2.

### *N. benthamiana* cell death assays

*A. tumefaciens* GV3101 (C58 (rifR) Ti pMP90 (pTiC58DT-DNA) (gentR) Nopaline (pSoup-tetR)) carrying wild-type or mutated Pik-1, Pik-2 were resuspended in agroinfiltration media (10 mM MgCl_2_, 10 mM 2-(N-morpholine)-ethanesulfonic acid (MES), pH 5.6) supplemented with 150 µM acetosyringone. Given combinations of Pik-1 and Pik-2 constructs were mixed at OD_600_ 0.4 for each construct. *A. tumefaciens* GV3101 carrying AVR-Pik effectors or mCherry were added to each experiment at OD_600_ 0.6. Each infiltration had additional *A. tumefaciens* GV3101 (C58 (rifR) Ti pMP90 (pTiC58DT-DNA) (gentR) Nopaline (pSoup-tetR)) carrying P19 at OD_600_ 0.1. Leaves of 4-weeks-old *N. benthamiana* were infiltrated using a needleless syringe. Leaves were collected at 5 dpi to measure UV autofluorescence as proxy for cell death as reported previously (De la Concepcion et al., 2019; De la Concepcion et al., 2018; Maqbool et al., 2015).

### Cell death scoring: UV autofluorescence

Detached leaves were imaged at 5 dpi from the abaxial side of the leaves for UV fluorescence images. Photos were taken using a Nikon D4 camera with a 60 mm macro lens, ISO set 1600 and exposure ∼10 secs at F14. The filter is a Kodak Wratten No.8 and white balance is set to 6250 degrees Kelvin. Blak-Ray® longwave (365nm) B-100AP spotlight lamps are moved around the subject during the exposure to give an even illumination. Images shown are representative of three independent experiments, with internal repeats. The cell death index used for scoring is as presented previously (Maqbool et al., 2015). Dotplots were generated using R v3.4.3 (https://www.r-project.org/) and the graphic package ggplot2 (Wickham, 2016). The size of the centre dot at each cell death value is directly proportional to the number of replicates in the sample with that score. All individual data points are represented as dots.

## Acknowledgments

We thank present and former members of the Banfield and Kamoun laboratories for discussions that have shaped this manuscript, and colleagues at Iwate Biotechnology Research Center for stimulating discussions on NLR biology. We specially thank Dr. Cristina Barragan and Dr. Adam Bentham for critical reading of the manuscript. We also thank Andrew Davies and Phil Robinson from JIC Scientific Photography for the UV pictures of the cell death assays. This work was supported by the Biotechnology and Biological Sciences Research Council (BBSRC, UK, grant BB/012574, BBS/E/J/000PR9795), the BBSRC Doctoral Training Partnership at Norwich Research Park (grant: BB/M011216/1, project reference 1771322), the European Research Council (proposal 743165), the John Innes Foundation, the Gatsby Charitable Foundation, the European Commission through the Erasmus+ programme, and JSPS Grant 20H05681.

---

**Figure S1.**
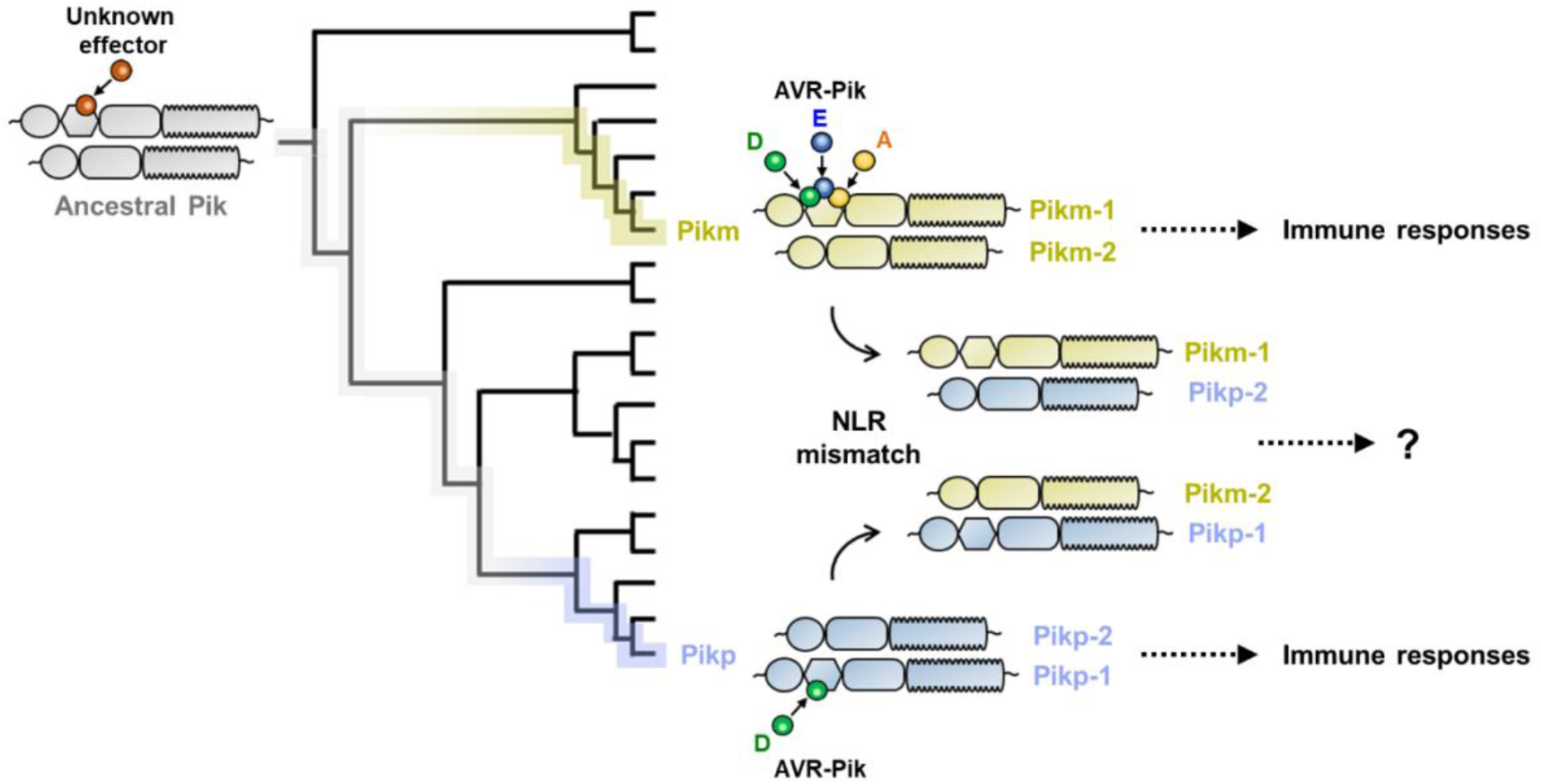
Schematic representation of the hypothesis tested in this study. Sensor NLR alleles Pikp-1 and Pikm-1 convergently evolved to bind *M. oryzae* AVR-Pik effectors, triggering immune responses together with their corresponding NLR pair. We tested sensor/helper specificity in Pikp and Pikm pairs by mismatching allelic receptor and measuring immune response outcomes.

**Figure S2.**
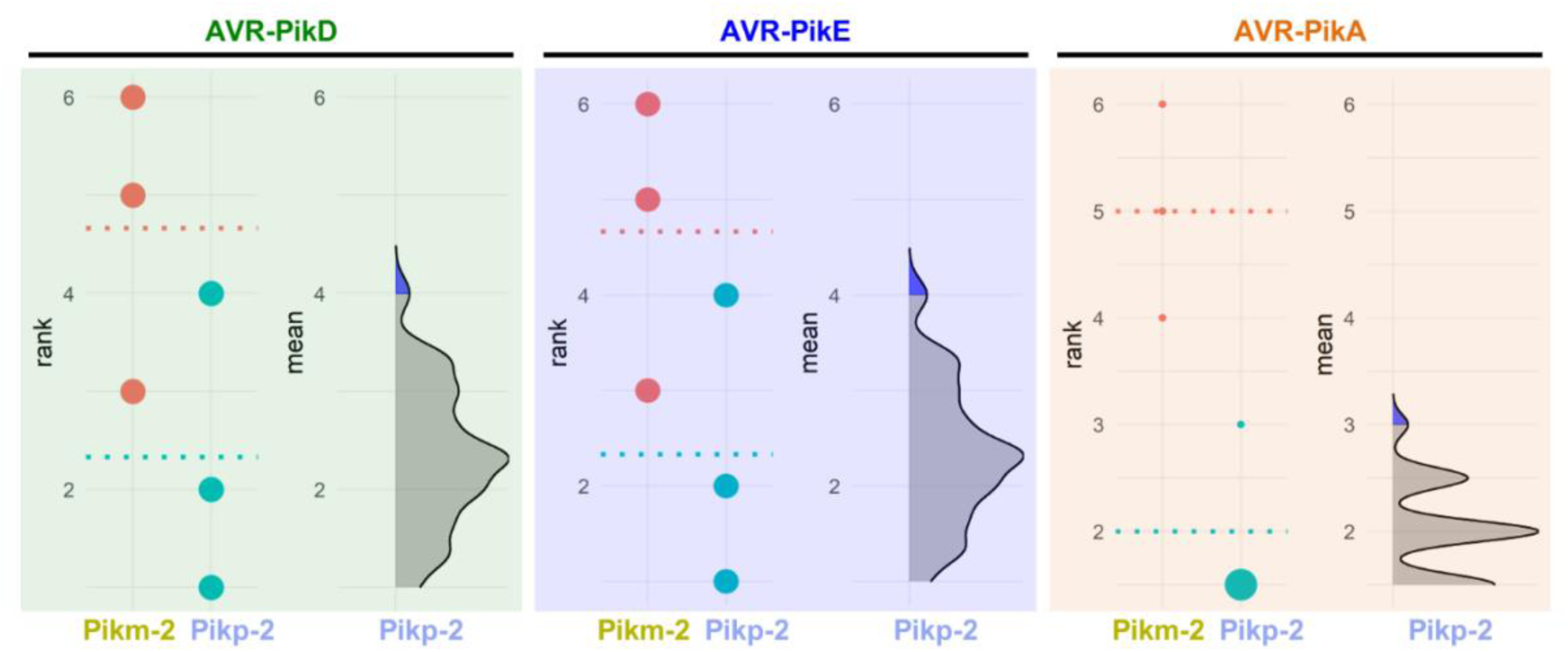
Estimation graphics for comparison of cell death mediated by Pikm-1 when co-expressed with Pikm-2 or Pikp-2. Statistical analysis by estimation methods of the cell death assay for Pikm-1 co-expressed with Pikm-2 or Pikp-2 and AVR-PikD, AVR-PikE or AVR-PikA. For each effector, the panel on the left represents the ranked data (dots) for each NLR, and their corresponding mean (dotted line). The size of the dots is proportional to the number of observations with that specific value. The panel on the right shows the distribution of 1000 bootstrap sample rank means for Pikm-1 paired with Pikp-2. The blue areas represent the 0.025 and 0.975 percentiles of the distribution. Pikm-1 mediated responses with Pikm-2 or Pikp-2 are considered significantly different if the Pikm-2 rank mean (dotted line, left panel) falls beyond the blue regions of the Pikp-2 mean distribution.

**Figure S3.**
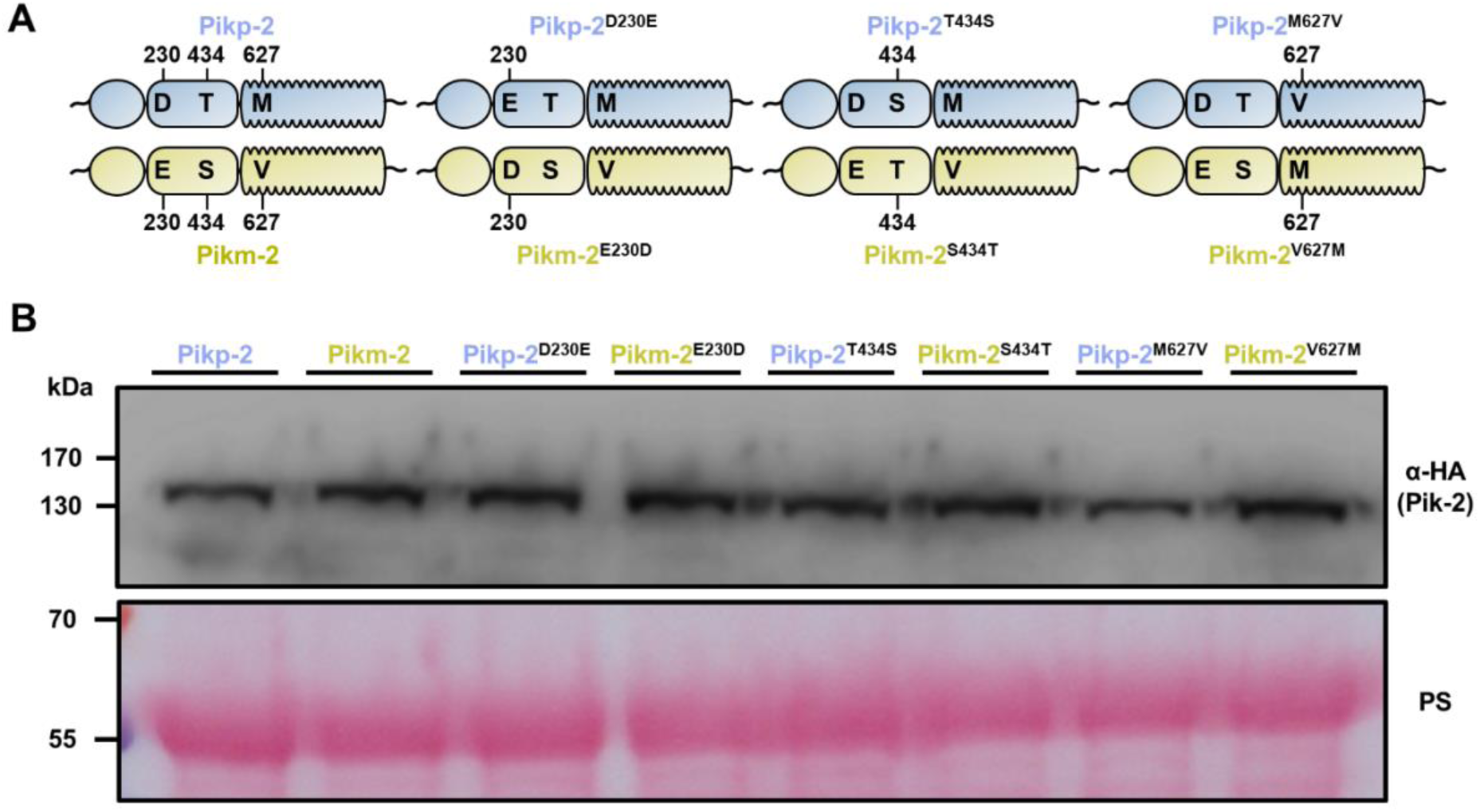
The Pik-2 alleles and mutants have similar levels of protein accumulation in planta. **(A)** Schematic representations of polymorphism distribution in the Pik-2 allelic NLRs and their mutants. Polymorphic sites are numbered. **(B)** Western blots showing accumulation of wild-type Pikp-2 and Pikm-2 and point mutants. C-terminally 6*×*HA tagged Pik-2 proteins were transiently expressed *N. benthamiana*. Total protein extracts were probed with α-HA antisera. Total protein loading is shown by Ponceau staining (PS).

**Figure S4.**
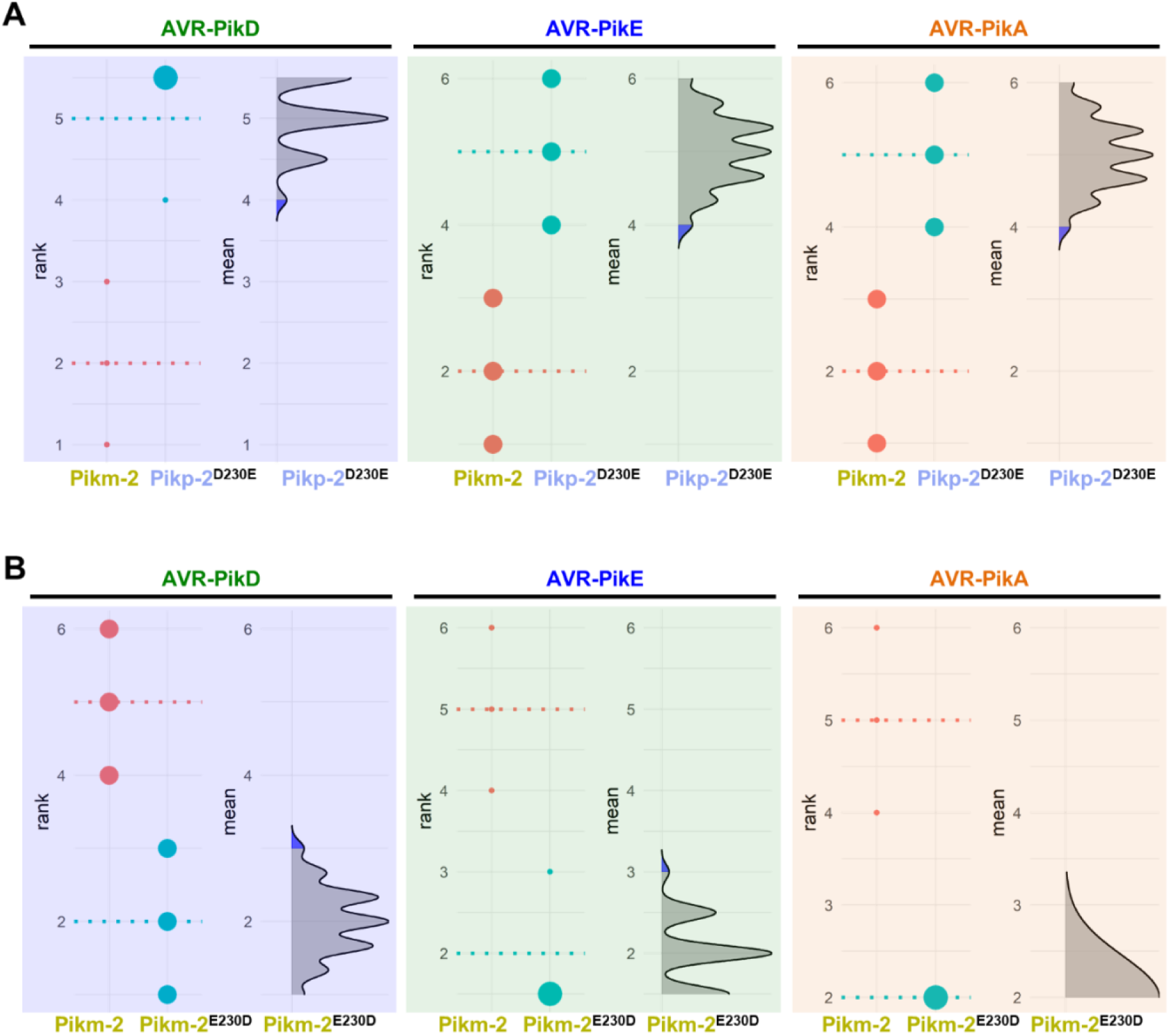
Estimation graphics for comparison of cell death mediated by Pikm-1 when co-expressed with the Pikm-2 or Pik-2 mutants in the polymorphic position 230. Statistical analysis by estimation methods of the cell death assay for Pikm-1 co-expressed with **(A)** Pikp-2 Asp230Glu or **(B)** Pikm-2 Glu230Asp and AVR-PikD, AVR-PikE or AVR-PikA, compared with wild-type Pikm-2. For each effector, the panel on the left represents the ranked data (dots) for each NLR, and their corresponding mean (dotted line). The size of the dots is proportional to the number of observations with that specific value. The panel on the right shows the distribution of 1000 bootstrap sample rank means for Pikm-1 paired with a Pik-2 mutant. The blue areas represent the 0.025 and 0.975 percentiles of the distribution. Pikm-1 mediated responses with Pikm-2 or Pik-2 mutant are considered significantly different if the Pikm-2 rank mean (dotted line, left panel) falls beyond the blue regions in the mean distribution of the Pik-2 mutants.

**Figure S5.**
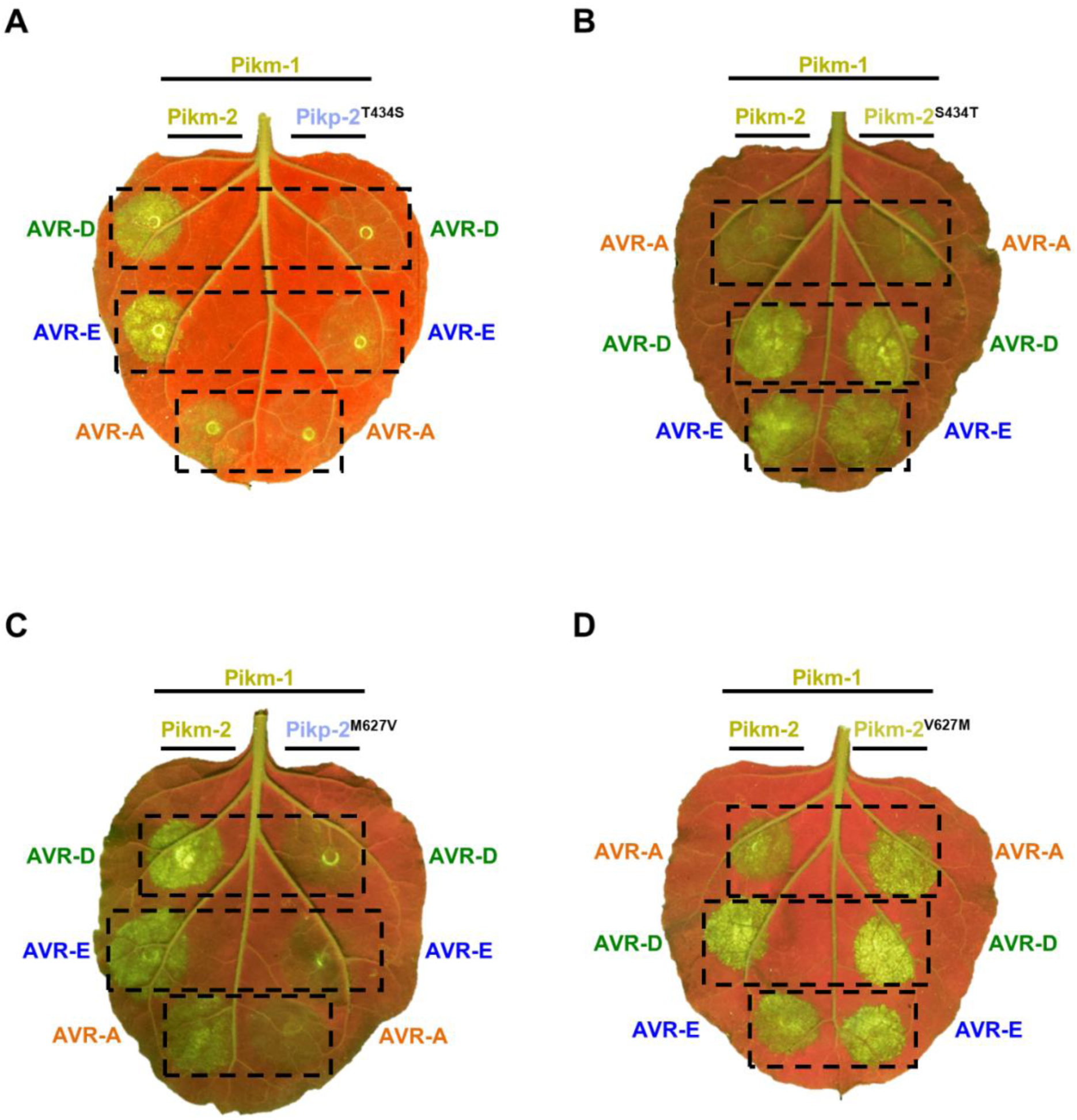
Representative images of cell death mediated by the Pik-2 mutants in response to the AVR-Pik effectors. Representative leaves depicting cell death mediated by Pik-2 mutants as autofluorescence under UV light. Pikm-1 was co-expressed with either **(A)** Pikp-2 Asp230Glu, **(B)** Pikm-2 Glu230Asp, **(C)** Pikp-2 Asp230Glu and **(D)** Pikm-2 Glu230Asp and AVR-PikD (AVR-D), AVR-PikE (AVR-E) or AVR-PikA (AVR-A). Side-by-side infiltrations with the Pikm NLR pair are highlighted with dashed boxes for comparison.

**Figure S6.**
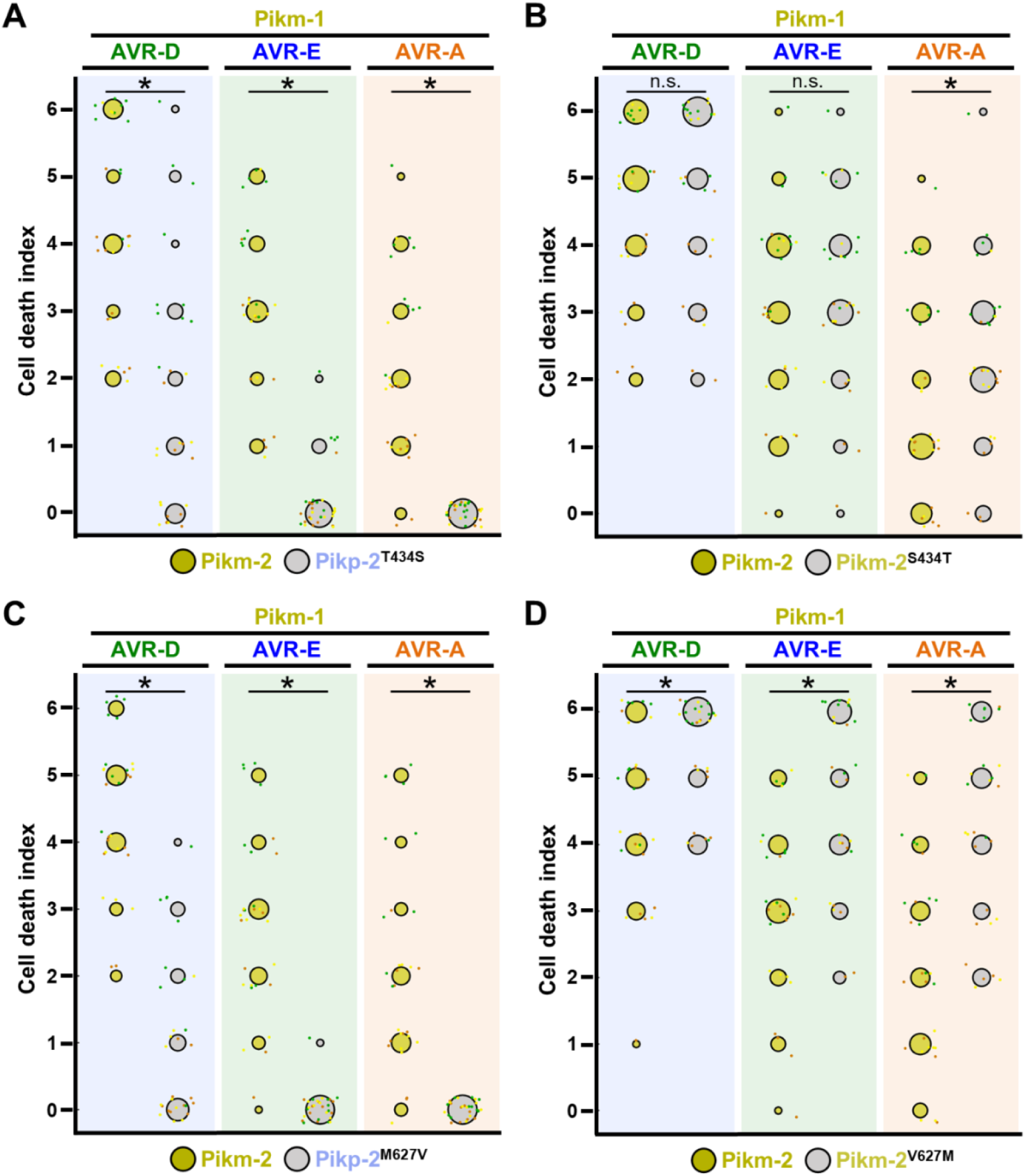
Quantification of cell death mediated by the Pik-2 mutants in response to AVR-Pik effectors. Cell death scoring is represented as dot plots comparing cell death triggered by the Pik-2 mutants **(A)** Pikp-2 Thr434Ser, **(B)** Pikm-2 Ser434Thr, **(C)** Pikp-2 Met627Val and **(D)** Pikm-2 Val627Met. The mutants were co-expressed with Pikm-1 and AVR-PikD, AVR-PikE or AVR-PikA. Pikm NLR pair was co-infiltrated for side-by-side comparison. The number of repeats was 30. For each sample, all the data points are represented as dots with a distinct colour for each of the three biological replicates; these dots are jittered about the cell death score for visualisation purposes. The size of the central dot at each cell death value is proportional to the number of replicates of the sample with that score. Significant differences between relevant conditions are marked with an asterisk and the details of the statistical analysis are summarised in **Figure S7**.

**Figure S7.**
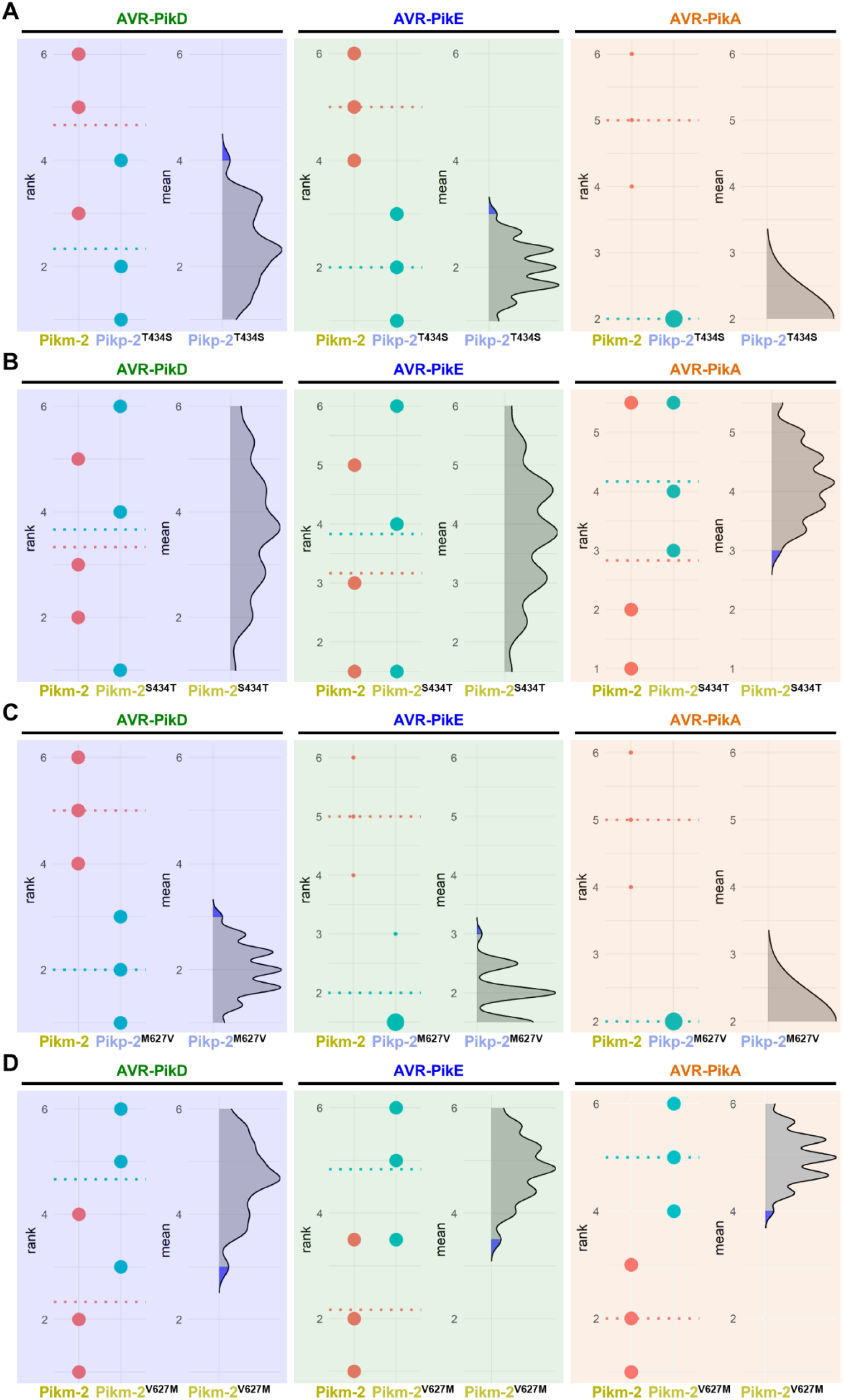
Estimation graphics for comparison of cell death mediated by Pikm-1 when co-expressed with Pikm-2 or Pik-2 mutants in polymorphic position 434 and 627. Statistical analysis by estimation methods of the cell death assay for Pikm-1 co-expressed with **(A)** Pikp-2 Thr434Ser, **(B)** Pikm-2 Ser434Thr, **(C)** Pikp-2 Met627Val or **(D)** Pikm-2 Val627Met and AVR-PikD, AVR-PikE or AVR-PikA, compared with wild-type Pikm-2. For each effector, the panel on the left represents the ranked data (dots) for each NLR, and their corresponding mean (dotted line). The size of the dots is proportional to the number of observations with that specific value. The panel on the right shows the distribution of 1000 bootstrap sample rank means for Pikm-1 paired with a Pik-2 mutant. The blue areas represent the 0.025 and 0.975 percentiles of the distribution. Pikm-1 mediated responses with Pikm-2 or Pik-2 mutant are considered significantly different if the Pikm-2 rank mean (dotted line, left panel) falls beyond the blue regions in the mean distribution of the Pik-2 mutants.

**Figure S8.**
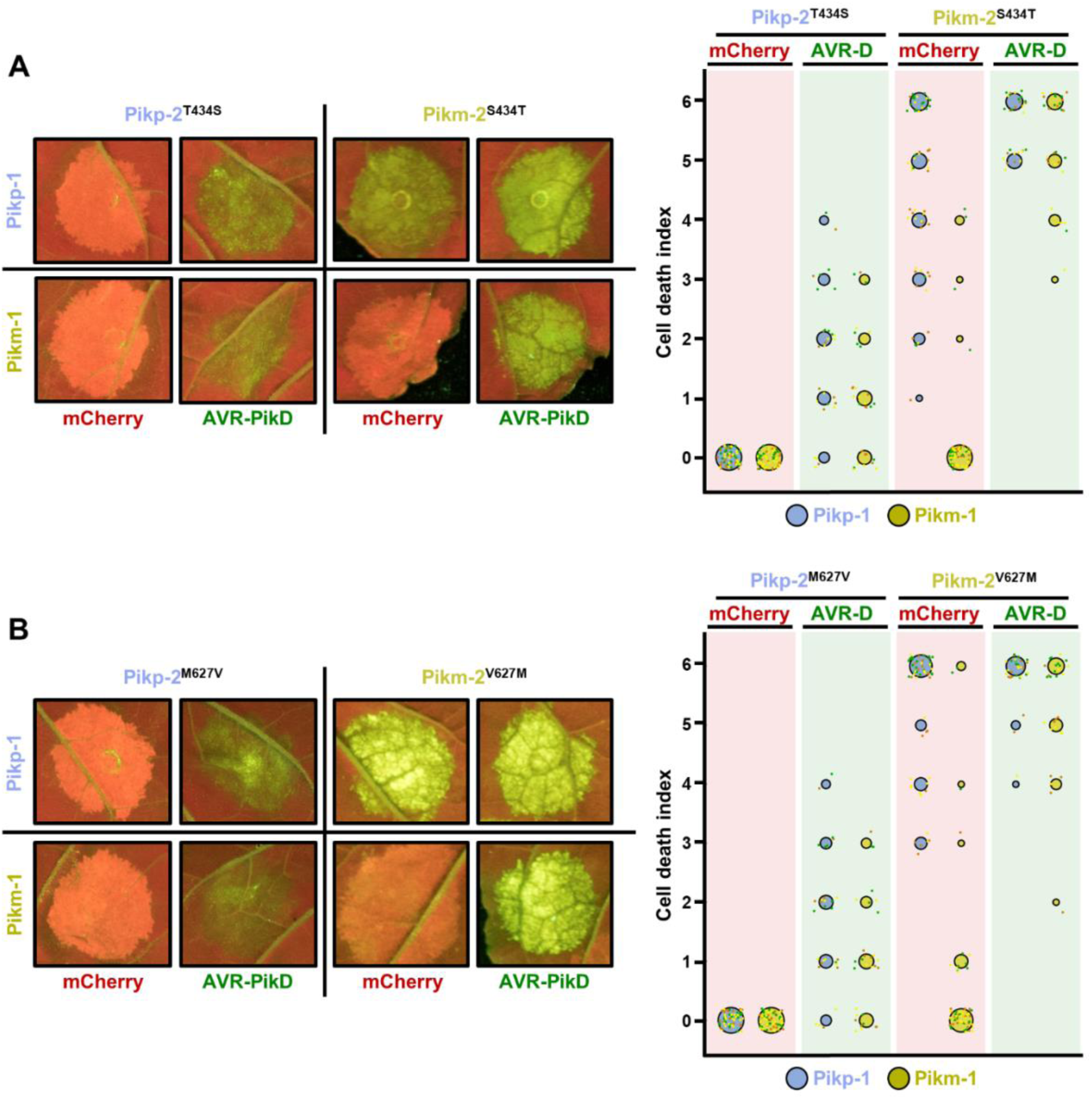
The Pik-2 polymorphisms at position 434 and 627 do not alter constitutive cell death. Representative leaf spot images and scoring of cell death mediated by Pik-2 as autofluorescence under UV-light. Cell death scoring is represented as dot plots comparing cell death triggered by Pik-2 mutants at polymorphic positions **(A)** 434 and **(B)** 627. Pik-2 mutants were co-expressed with Pikp-1 (blue dots) or Pikm-1 (yellow dots) together with mCherry (red panel) or AVR-PikD (green panel). The number of repeats was 60 and 30 for the spots co-infiltrated with mCherry and AVR-PikD, respectively. For each sample, all the data points are represented as dots with a distinct colour for each of the three biological replicates; these dots are jittered about the cell death score for visualisation purposes. The size of the central dot at each cell death value is proportional to the number of replicates of the sample with that score.

**Figure S9.**
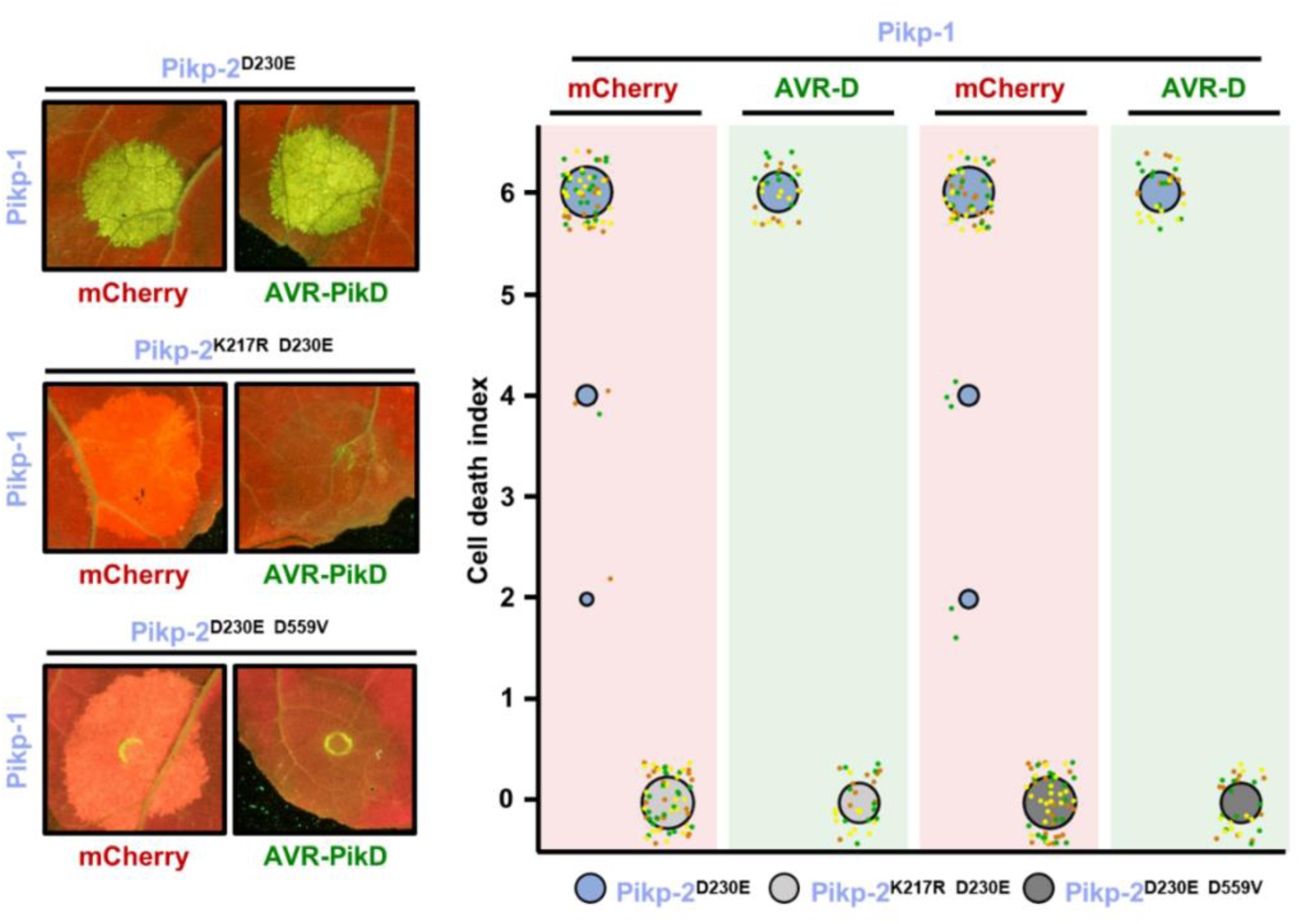
Pikp-2 Asp230Glu autoactivation is dependent on P-loop and MHD motifs. Representative leaf spot images and scoring of Pikm-2 mediated cell death as autofluorescence under UV-light. Cell death scoring is represented as dot plots comparing cell death triggered by Pikp-2 Asp230Glu mutant and its versions mutated in P-loop (Lys217Arg) and MHD (Asp559Val) motifs. Pik-2 mutants were co-expressed with Pikp-1 and mCherry (red panel) or AVR-PikD (green panel). The number of repeats was 60 and 30 for the spots co-infiltrated with mCherry and AVR-PikD, respectively. For each sample, all the data points are represented as dots with a distinct colour for each of the three biological replicates; these dots are jittered about the cell death score for visualisation purposes. The size of the central dot at each cell death value is proportional to the number of replicates of the sample with that score.

**Figure S10.**
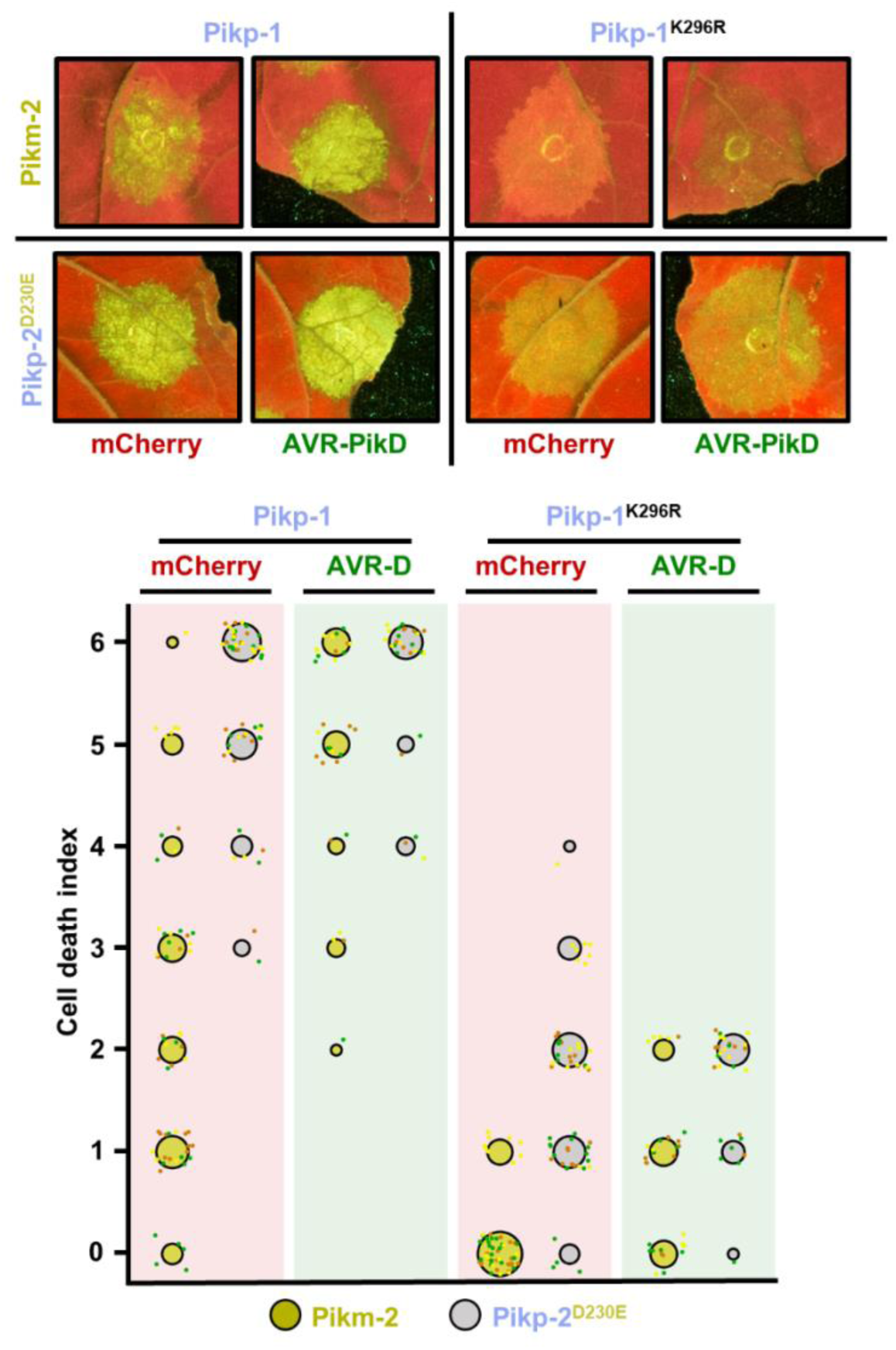
The Pik-1 P-loop motif is important but not essential for Pik-mediated cell death. Representative leaf spot images and scoring of Pik-mediated cell death as autofluorescence under UV-light. Cell death scoring is represented as dot plots comparing cell death triggered by Pikm-2 or Pikp-2 Asp230Glu in the presence of wild-type Pikp-1 or a version mutated in the P-loop motif (Lys296Arg). The different NLR pair combinations were co-infiltrated with mCherry (red panel) or AVR-PikD (green panel). The number of repeats was 60 and 30 for the spots co-infiltrated with mCherry and AVR-PikD, respectively. For each sample, all the data points are represented as dots with a distinct colour for each of the three biological replicates; these dots are jittered about the cell death score for visualisation purposes. The size of the central dot at each cell death value is proportional to the number of replicates of the sample with that score.

**Figure S11.**
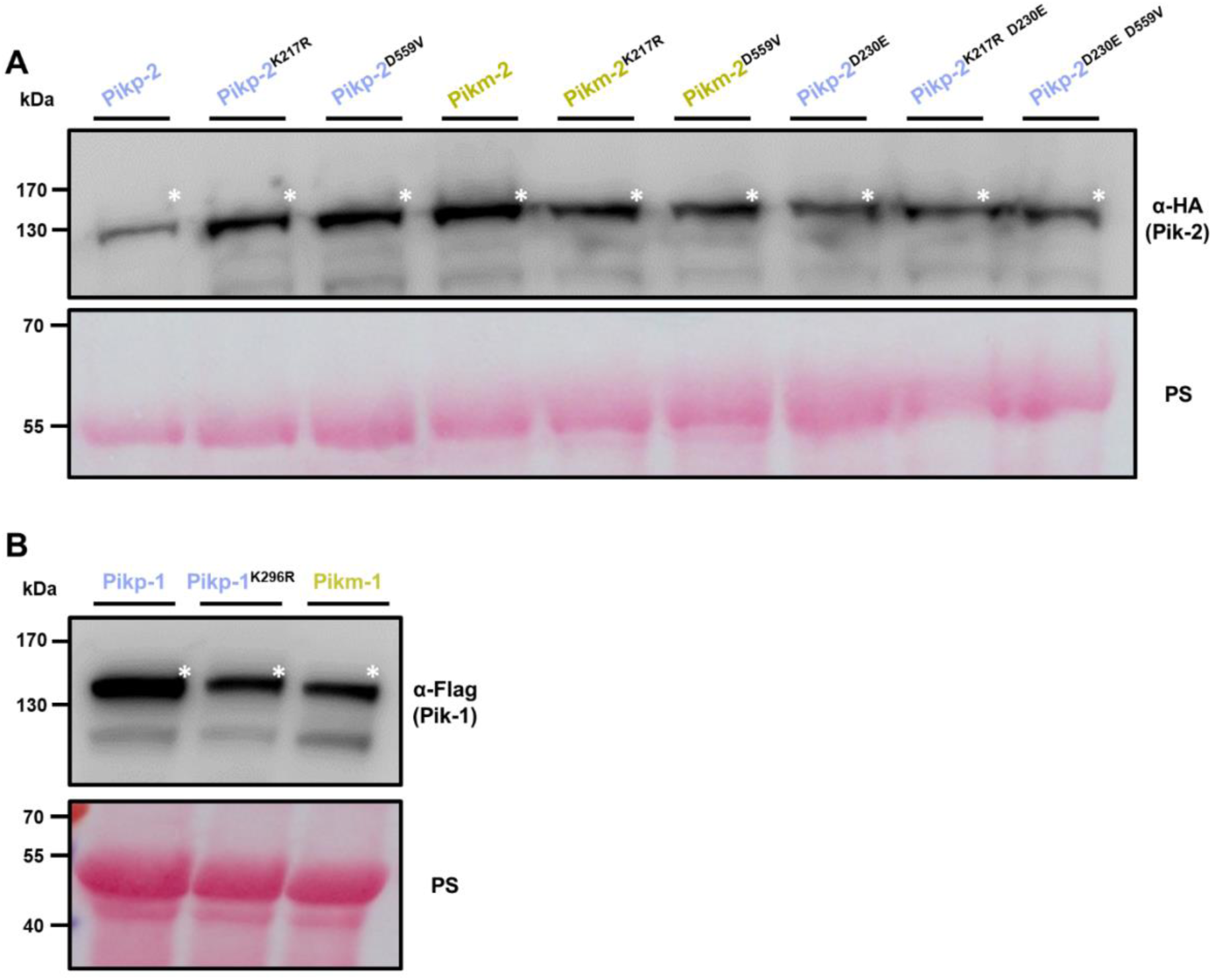
Mutations in P-loop and MHD motifs do not affect protein accumulation. Western blots showing accumulation of **(A)** P-loop (Lys217R) and MHD (Asp559Val) mutants in the background of Pikp-2, Pikm-2 and Pikp-2 Asp230Glu. C-terminally 6*×*HA tagged Pik-2 mutants were transiently expressed *N. benthamiana*. C-terminally 6*×*HA tagged Pikp-2, Pikm-2 and Pikp-2 Asp230Glu are included as controls in each case. **(B)** Pikp-1 P-loop (Lys296R) mutant. C-terminally 6*×*His3*×*FLAG tagged Pikp-1 Lys296Arg mutant was transiently expressed *N. benthamiana*. C-terminally 6*×*His3*×*FLAG tagged wild-type Pikp-1 and Pikm-1 are included as controls (left and right, respectively). Total protein extracts were probed α-HA and α-FLAG antisera for A and B, respectively. Asterisks mark the band corresponding to the relevant protein. Total protein loading is shown by Ponceau staining (PS).

**Figure S12.**
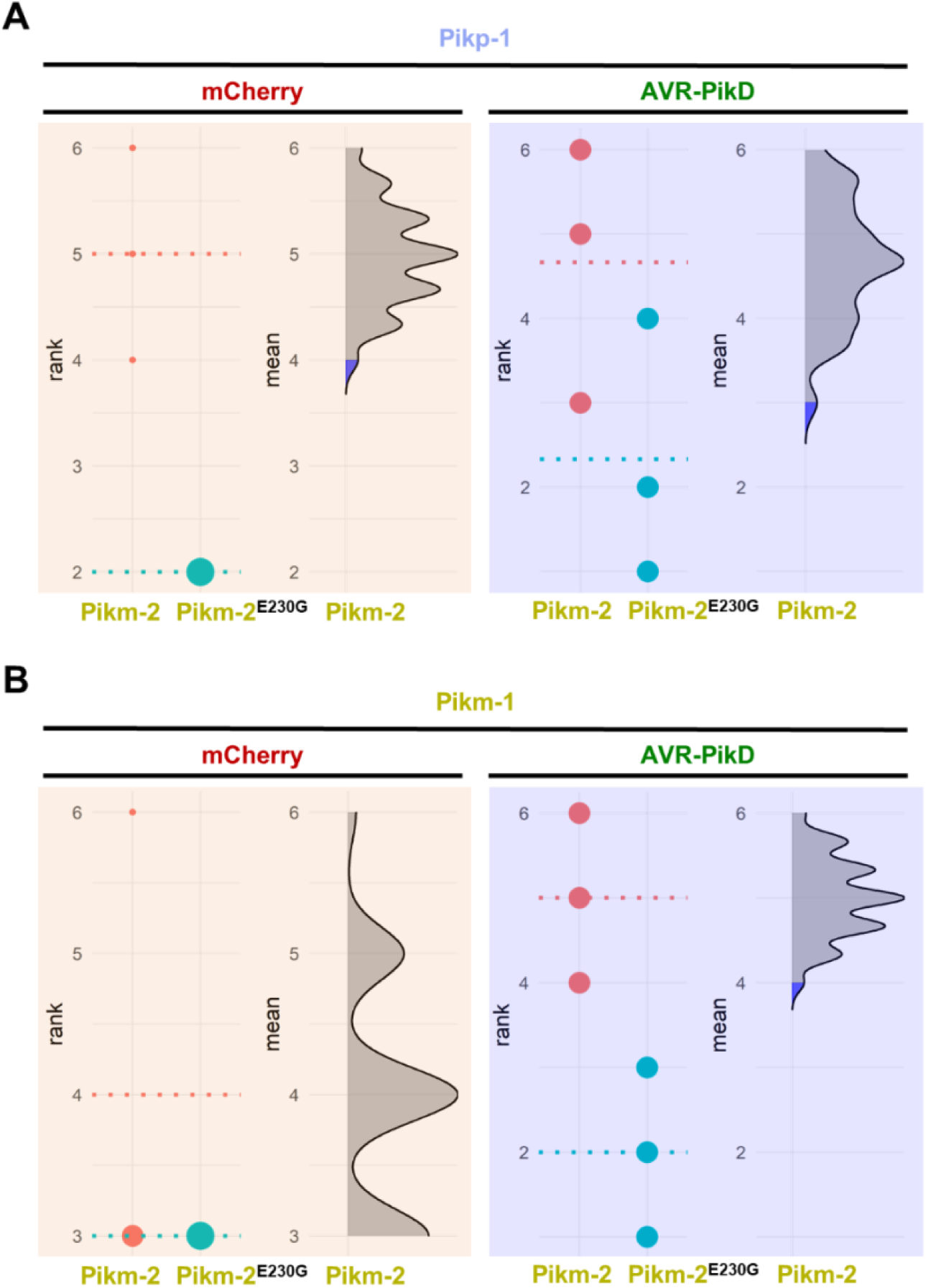
Estimation graphics for comparison of cell death mediated by Pikm-2 or Pikm-2 Glu230Gly. Statistical analysis by estimation methods of the cell-death assay for **(A)** Pikp-1 or **(B)** Pikm-1 co-expressed with Pikm-2 Glu230Gly and mCherry or AVR-PikD, compared with wild-type Pikm-2. The panel on the left represents the ranked data (dots) for each NLR, and their corresponding mean (dotted line). The size of the dots is proportional to the number of observations with that specific value. The panel on the right shows the distribution of 1000 bootstrap sample rank means for Pik-1 paired with Pikm-2. The blue areas represent the 0.025 and 0.975 percentiles of the distribution. Pikm-2 Glu230Gly mediated responses are considered significantly different if the Pikm-2 rank mean (dotted line, left panel) falls beyond the blue regions of the Pikm-2 Glu230Gly mean distribution.

**Figure S13.**
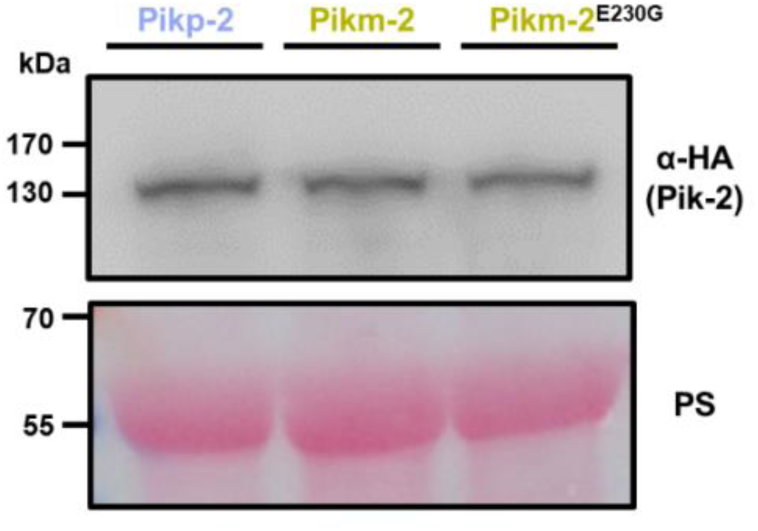
Glu230Gly mutation does not affect Pik-2 protein accumulation. Western blots showing accumulation of Pikm-2 Glu230Gly. C-terminally 6*×*HA tagged Pikm-2 Glu230Gly mutant was transiently expressed *N. benthamiana*. C-terminally 6*×*HA tagged Pikp-2 and Pikm-2 alleles are included as controls. Total protein extracts were probed α-HA antisera. Total protein loading is shown by Ponceau staining (PS).

**Figure S14.**
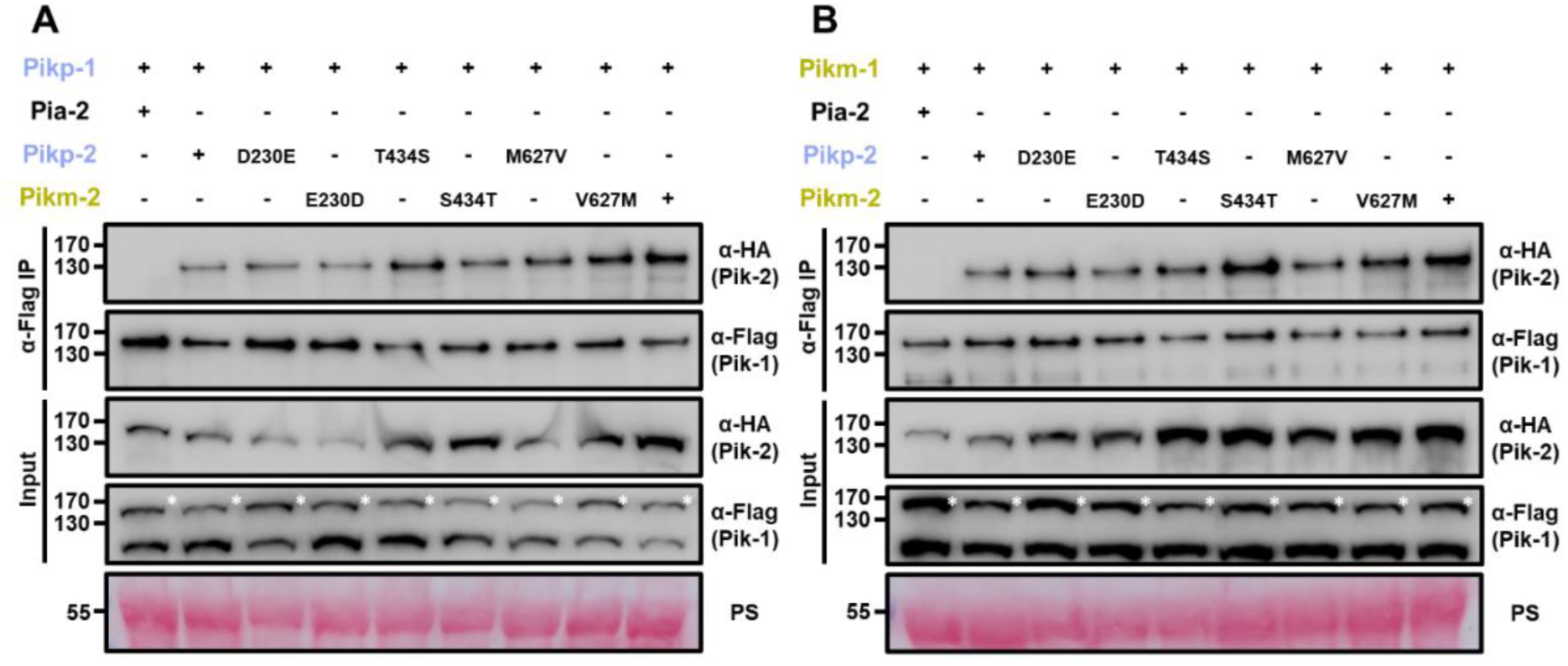
Pik-2 mutants associate with Pik-1 in planta. Co-immunoprecipitation of full length Pikp-1 **(A)** or Pikm-1 **(B)** with each Pik-2 mutant in polymorphic sites. C-terminally 6*×*HA tagged Pik-2 NLR mutants were transiently co-expressed with Pik-1:6*×*His3*×*FLAG in *N. benthamiana*. Immunoprecipitates obtained with anti-FLAG antiserum, and total protein extracts, were probed with appropriate antisera. Co-expression with C-terminally tagged 6*×*HA Pia-2 NLR is included as negative control. Asterisks mark the band corresponding to Pikp-1. Total protein loading is shown by Ponceau staining (PS).

**Figure S15.**
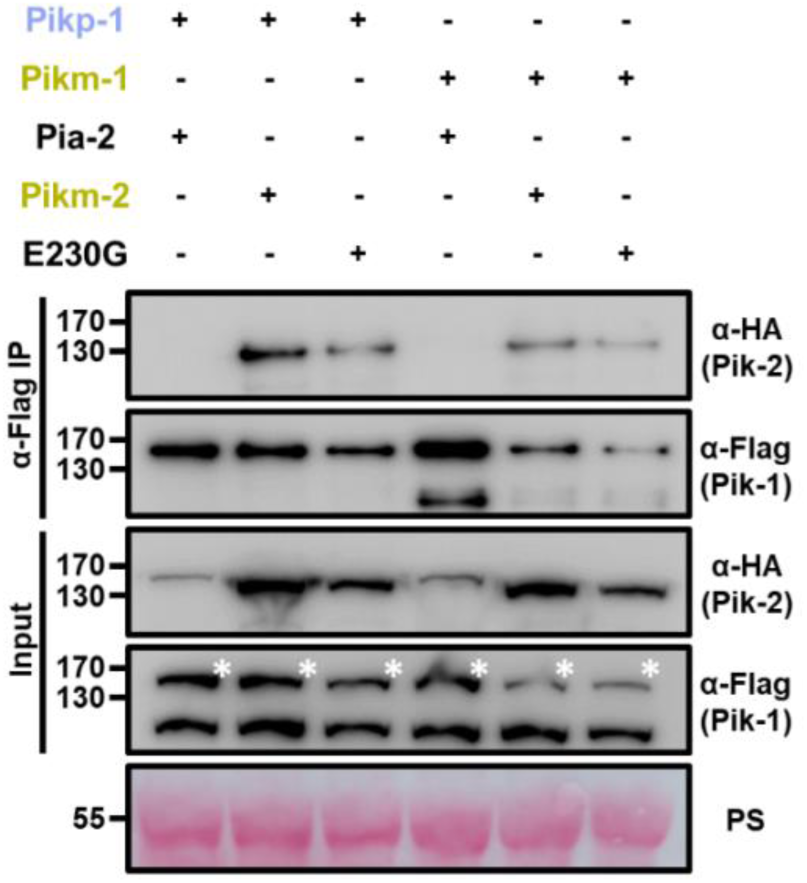
Reversion to ancestral state in polymorphism 230 does not abrogate association with Pik-1 alleles. Co-immunoprecipitation of Pikm-2 Glu230Gly mutant with full length Pikp-1 and Pikm-1 alleles. C-terminally 6*×*HA tagged Pikm-2 Glu230Gly was transiently co-expressed with either Pikp-1:6*×*His3*×*FLAG or Pikm-1:6*×*His3*×*FLAG in *N. benthamiana*. Immunoprecipitates obtained with anti-FLAG antiserum, and total protein extracts, were probed with appropriate antisera. Co-expression with C-terminally tagged 6*×*HA Pia-2 NLR and wild-type Pikm-2 were included as negative and positive control, respectively. Asterisks mark the band corresponding to Pik-1. Total protein loading is shown by Ponceau staining (PS).

**Figure S16.**
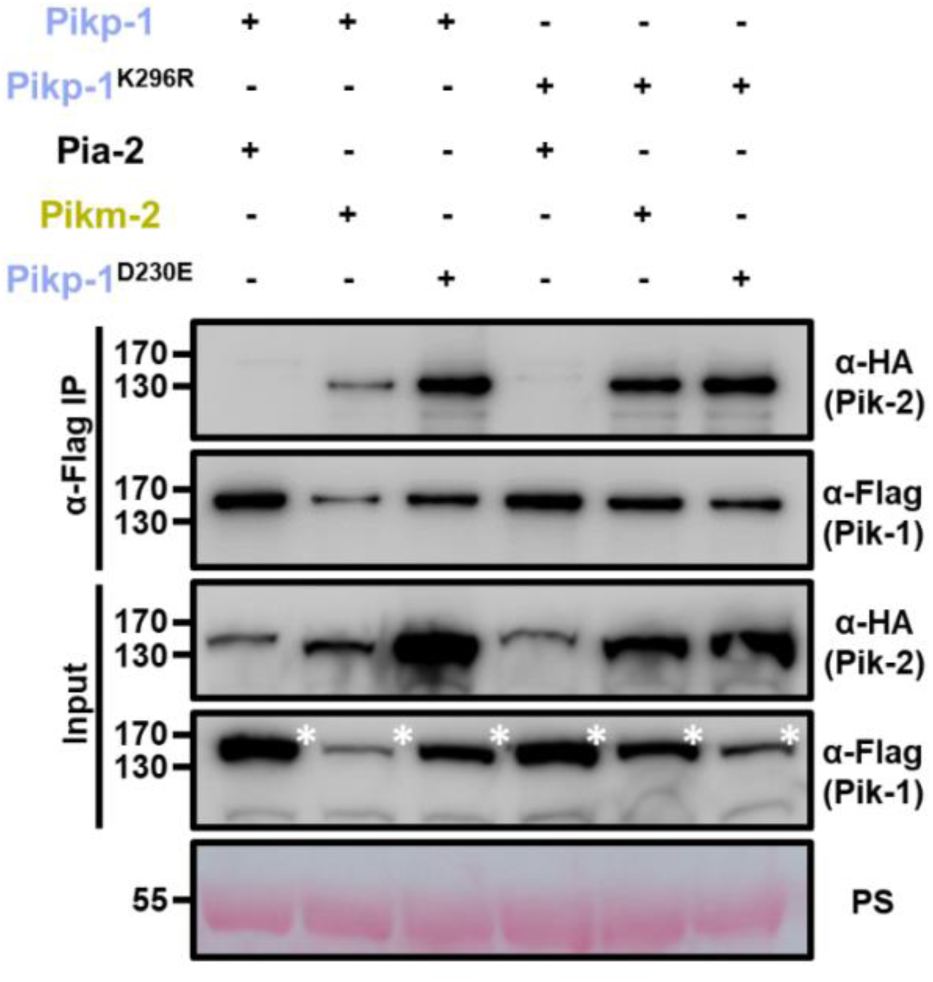
P-loop mutations does not affect Pik-1 association to Pik-2. Co-immunoprecipitation of Pikm-2 and Pikp-2 Asp230Glu with wild-type Pikp-1 and Pikp-1 P-loop mutant (Lys296Arg). C-terminally 6*×*HA tagged Pikm-2 and Pikp-2 Asp230Glu were transiently co-expressed with C-terminally 6*×*His3*×*FLAG tagged wild-type Pikp-1 or Pikp-1 Lys296Arg in *N. benthamiana*. Immunoprecipitates obtained with anti-FLAG antiserum, and total protein extracts, were probed with appropriate antisera. Co-expression with C-terminally tagged 6*×*HA Pia-2 NLR was included as negative control, respectively. Asterisks mark the band corresponding to Pik-1. Total protein loading is shown by Ponceau staining (PS).

**Figure S17.**
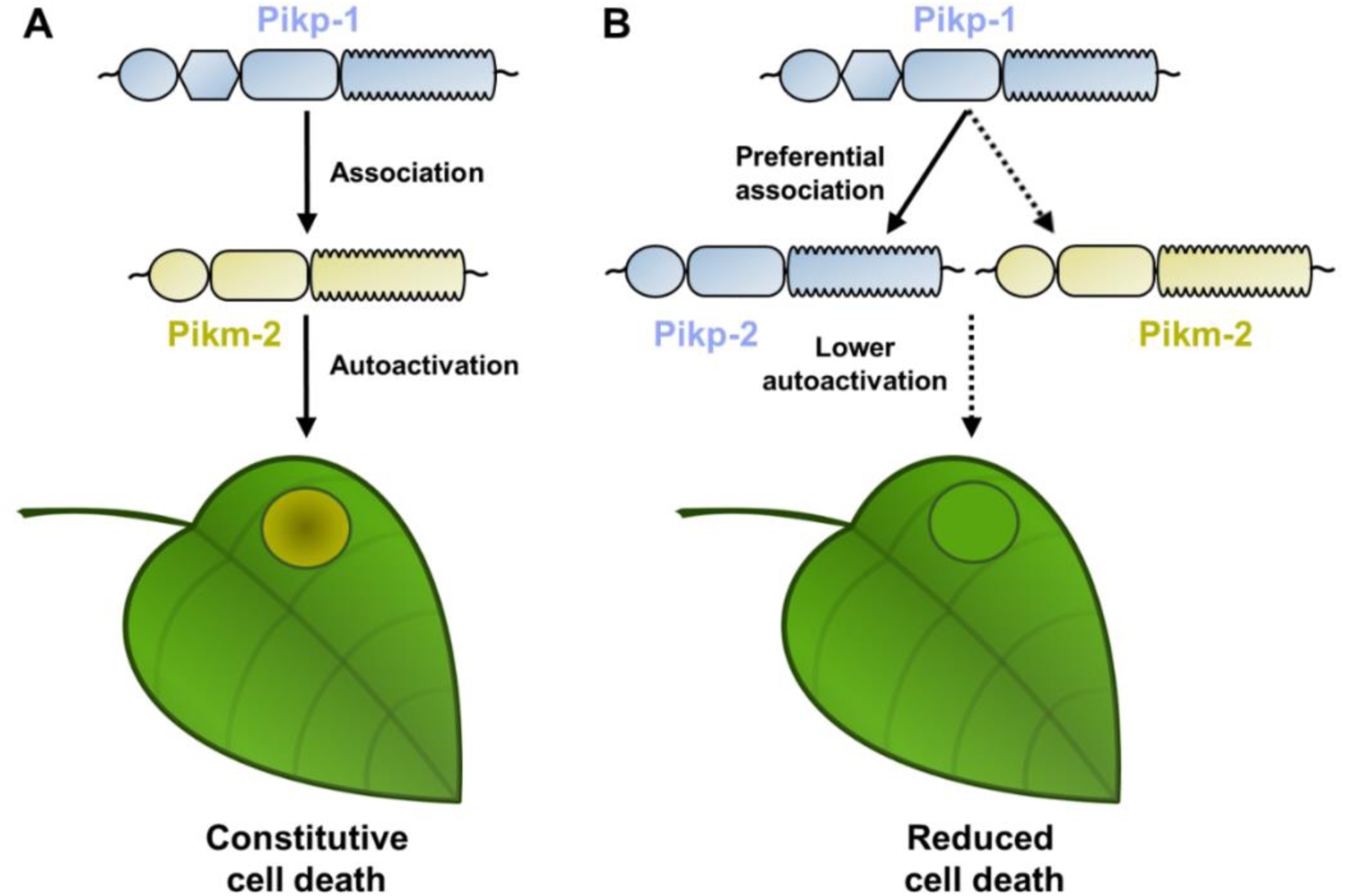
Schematic representations of Pik NLR competition assays. **(A)** When Pikp-1 (coloured in ice blue) is co-expressed with Pikm-2 (coloured in gold), both NLRs associate and trigger NLR activation that leads to constitutive cell death in *N. benthamiana*, depicted by the development of chlorotic and necrotic leaf tissue. **(B)** In a preferential association scenario, with both Pikp-2 and Pikm-2 present, Pikp-1 would associate with coevolved Pikp-2 instead of to Pikm-2 (depicted by the solid and dashed lines, respectively). This would reduce constitutive immune signalling and cell death.

**Figure S18.**
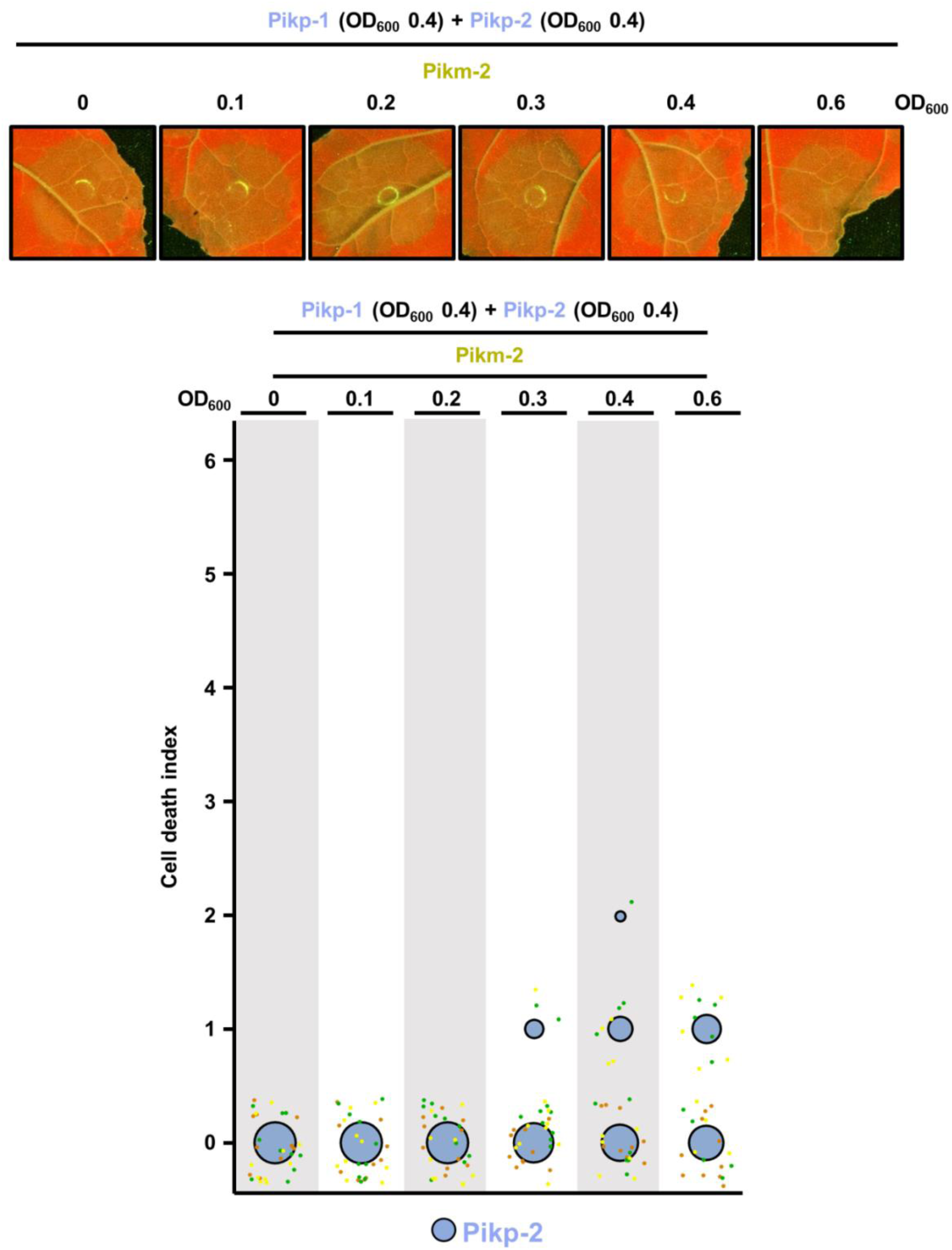
Pikp-2 suppresses constitutive cell death mediated by Pikm-2. Representative leaf spot images depicting Pikm-2 mediated cell death in the presence of Pikp-1 and Pikp-2 and increasing concentration of Pikm-2 as autofluorescence under UV-light. Scoring of the cell death assay is represented as dot plots. For each experiment, Pikp-1 and Pikp-2 were co-infiltrated at OD_600_ 0.4 each. Increasing concentrations of Pikm-2 were added to each experiment (from left to right: OD_600_ 0, 0.1, 0.2, 0.3, 0.4 and 0.6). A total of three biological replicates with 10 internal repeats each were performed for each experiment. For each sample, all the data points are represented as dots with a distinct colour for each of the three biological replicates; these dots are jittered about the cell death score for visualisation purposes. The size of the central dot at each cell death value is proportional to the number of replicates of the sample with that score.

